# Parallel adaptation in autopolyploid *Arabidopsis arenosa* is dominated by repeated recruitment of shared alleles

**DOI:** 10.1101/2021.01.15.426785

**Authors:** Veronika Konečná, Sian Bray, Jakub Vlček, Magdalena Bohutínská, Doubravka Požárová, Rimjhim Roy Choudhury, Anita Bollmann-Giolai, Paulina Flis, David E Salt, Christian Parisod, Levi Yant, Filip Kolář

## Abstract

Relative contributions of pre-existing vs *de novo* genomic variation to adaptation are poorly understood, especially in polyploid organisms, which maintain increased variation. We assess this in high resolution using autotetraploid *Arabidopsis arenosa*, which repeatedly adapted to toxic serpentine soils that exhibit skewed elemental profiles. Leveraging a fivefold replicated serpentine invasion, we assess selection on SNPs and structural variants (TEs) in 78 resequenced individuals and discovered substantial parallelism in candidate genes involved in ion homeostasis. We further modelled parallel selection and inferred repeated sweeps on a shared pool of variants in nearly all these loci, supporting theoretical expectations. A single, striking exception is represented by TWO PORE CHANNEL 1, which exhibits convergent evolution from independent *de novo* mutations at an identical, otherwise conserved site at the calcium channel selectivity gate. Taken together, this suggests that polyploid populations can rapidly adapt to environmental extremes, calling on both pre-existing variation and novel polymorphisms.

## Introduction

Rapid adaptation to novel environments is thought to be enhanced by the availability of genetic variation; however, the relative contribution of standing variation versus the role of novel mutation is a matter of debate^1,2^, especially in higher ploidy organisms. Whole genome duplication (WGD, leading to polyploidisation) is a major force underlying diversification across eukaryotic kingdoms, seen most clearly in plants^3–5^ with various effects on genetic variation^6–8^. While WGD is clearly associated with environmental change or stress^5,9^, the precise impact of WGD on adaptability is largely unknown in multicellular organisms, and there is virtually no work assessing the evolutionary sources of adaptive genetic variation in young polyploids. Work in autopolyploids, which clearly isolate effects of WGD from hybridisation (which is confounded in allopolyploids), indicates subtle genomic changes may follow WGD alone^8,10,11^, which raises the question of whence their adaptive value may originate.

Autopolyploidy is expected to alter the selection and adaptation process in many ways, but a dearth of empirical data prevents synthetic evaluation. Besides immediate phenotypic^12–14^ and genomic^10,11^ changes following WGD, theory is unsettled regarding how adaptation proceeds as the autopolyploid lineage diversifies and adapts to novel challenges. On the one hand, autopolyploids can better mask deleterious alleles and accumulate cryptic allelic diversity^7^. In addition, the number of mutational targets is multiplied in autopolyploids, meaning that new alleles are introduced more quickly^6,15,16^. This could lead to higher rates of adaptation^6,17^. On the other hand, changes in allele frequency take longer in autopolyploids, which may retard adaptation^6,18^, particularly for *de novo* mutations, which emerge in a population at initially low frequencies^19^. Recent advances in theory and simulations suggest potential solutions to this controversy. Polyploidy may promote adaptation under scenarios of rapid environmental change (e.g. colonisation of challenging habitats) when selection is strong and originally neutral or mildly deleterious alleles standing in highly variable polyploid populations may become beneficial^5,20^. However, empirical evidence supporting this scenario is fragmentary. There is broad correlative evidence that polyploids are better colonizers of areas experiencing environmental flux (e.g. the Arctic^21,22^, stressful habitats^8,23,24^, and heterogeneous environments^25^). However, the genomic basis of such polyploid adaptability – and whether their primary source of adaptive alleles is high diversity (standing variation) or large mutational target size (*de novo* mutations) – remains unknown.

Here we deconstruct the sources of adaptive variation in natural autotetraploid *Arabidopsis arenosa* populations repeatedly facing one of the greatest environmental challenges for plant life – naturally toxic serpentine soils. Serpentines occur as islands in the landscape with no intermediate habitats and are defined by peculiar elemental contents (highly skewed Ca:Mg ratio and elevated heavy metals such as Cr, Co, and Ni), that may be further combined with low nutrient availability and propensity for drought^26^. *Arabidopsis arenosa* is a well characterized, natural diploid-autotetraploid species with large outcrossing populations^27^. The widespread autotetraploids, which originated from a single diploid lineage ∼19-31k generations ago^8^, harbour increased adaptive diversity genome-wide^8^ and currently occupy a broader ecological niche than their diploid sisters^28^, including serpentine outcrops^29^, railway lines^30,31^, and contaminated mine tailings^32,33^. This makes *A. arenosa* a promising model for empirical inquiries of adaptation in autopolyploids^8,27^. As a proof-of-concept, selective ion uptake phenotypes and a polygenic basis for serpentine adaptation have been suggested from a single *A. arenosa* serpentine population^29^. However, limited sampling left unknown whether the same genes are generally (re)used and what is the evolutionary source of the selected alleles, i.e. leaving unknown the evolutionary dynamics and mechanism underlying the striking adaptations.

To address these questions, we first sample five serpentine/non-serpentine population pairs of autotetraploid *A. arenosa* and demonstrate rapid parallel adaptation by combining demographic analysis and reciprocal transplant experiments. Taking advantage of the power of this five-fold replicated natural selection experiment, we identify candidate adaptive loci from population resequencing data and find substantial parallelism underlying serpentine adaptation. We then model parallel selection using a designated framework and statistically infer the evolutionary sources of parallel adaptive variation for all candidate loci. We ask specifically: (1) Does *A. arenosa* adapt dominantly via repeated sampling from the large pool of shared variation that is expected to be maintained in autopolyploids? and (2) Is repeated adaptation from novel mutations feasible in autotetraploid populations? In line with theory, we found that shared variation is the vastly prevalent source of adaptive variants in serpentine *A. arenosa*. However, we also discovered an exceptional locus exhibiting footprints of selection on alleles originating from two distinct *de novo* mutations. In line with the latter hypothesis, this demonstrates that the rapid selection of novel alleles is still feasible in autopolyploids, indicating broad evolutionary flexibility of lineages with doubled genomes.

## Results

### Parallel serpentine adaptation

First, we tested for independent colonisation of each serpentine site by different local *A. arenosa* populations. To do this, we resequenced five pairs of geographically proximate serpentine (S) and non-serpentine (N) populations covering all known serpentine sites occupied by the species to date (8 individuals per population on average, mean sequencing depth 21×; Fig 1a, Fig S1, Dataset S1, and Table S1, S2, S3). Phylogenetic, ordination and Bayesian analyses based on nearly-neutral fourfold-degenerate sites demonstrated grouping of populations by spatial proximity, not by substrate (Fig 1b, Fig S1b, d). Independent colonization of each serpentine site was confirmed by coalescent simulations in which the scenario involving parallel local colonisation of each serpentine site was more likely than the complementary scenario assuming monophyly of the substrate type (S vs. N), followed by admixture. This was consistent over all possible pairwise iterations of population pairs (n=10) (Fig 1c, Fig S2). The very low differentiation between serpentine and sister non-serpentine populations and consistently low population split times (Table 1) indicate very recent, postglacial serpentine invasions. There is no evidence of bottleneck associated with colonization, as serpentine and non-serpentine populations exhibited similar nucleotide diversity and Tajima’s D values (Table 1).

**Table 1.** Between population divergence and within population diversity of the five investigated serpentine / non-serpentine population pairs.

| Pop. pair | Divergence (Generations)<sup>1</sup> | Pairwise F<sub>ST</sub> | Nucleotide diversity<sup>2</sup> | Tajima's D<sup>2</sup> |
| --- | --- | --- | --- | --- |
| S1-N1 | (774) 4,317 (6,284) | 0.069 | 0.0292 / 0.0276 | 0.266 / 0.485 |
| S2-N2 | (500) 2,690 (3,826) | 0.029 | 0.0306 / 0.0285 | 0.131 / 0.372 |
| S3-N3 | (794) 3,912 (6,316) | 0.085 | 0.0307 / 0.0287 | -0.046 / 0.350 |
| S4-N4 | (812) 2,918 (3,542) | 0.057 | 0.0304 / 0.0297 | 0.132 / 0.177 |
| S5-N5 | (546) 3,539 (4,869) | 0.047 | 0.0292 / 0.0296 | 0.267 / 0.089 |

<sup>1</sup>Divergence between proximal S-N populations. Mean and 95% confidence intervals inferred by bootstrapping, estimated by coalescent simulations. Assuming two-year generation time<sup>34</sup>, all estimates indicate recent postglacial divergence.
<sup>2</sup>Genome-wide nucleotide diversity ( $\pi$ ) and Tajima's D of each S / N population.

**Fig 1.**
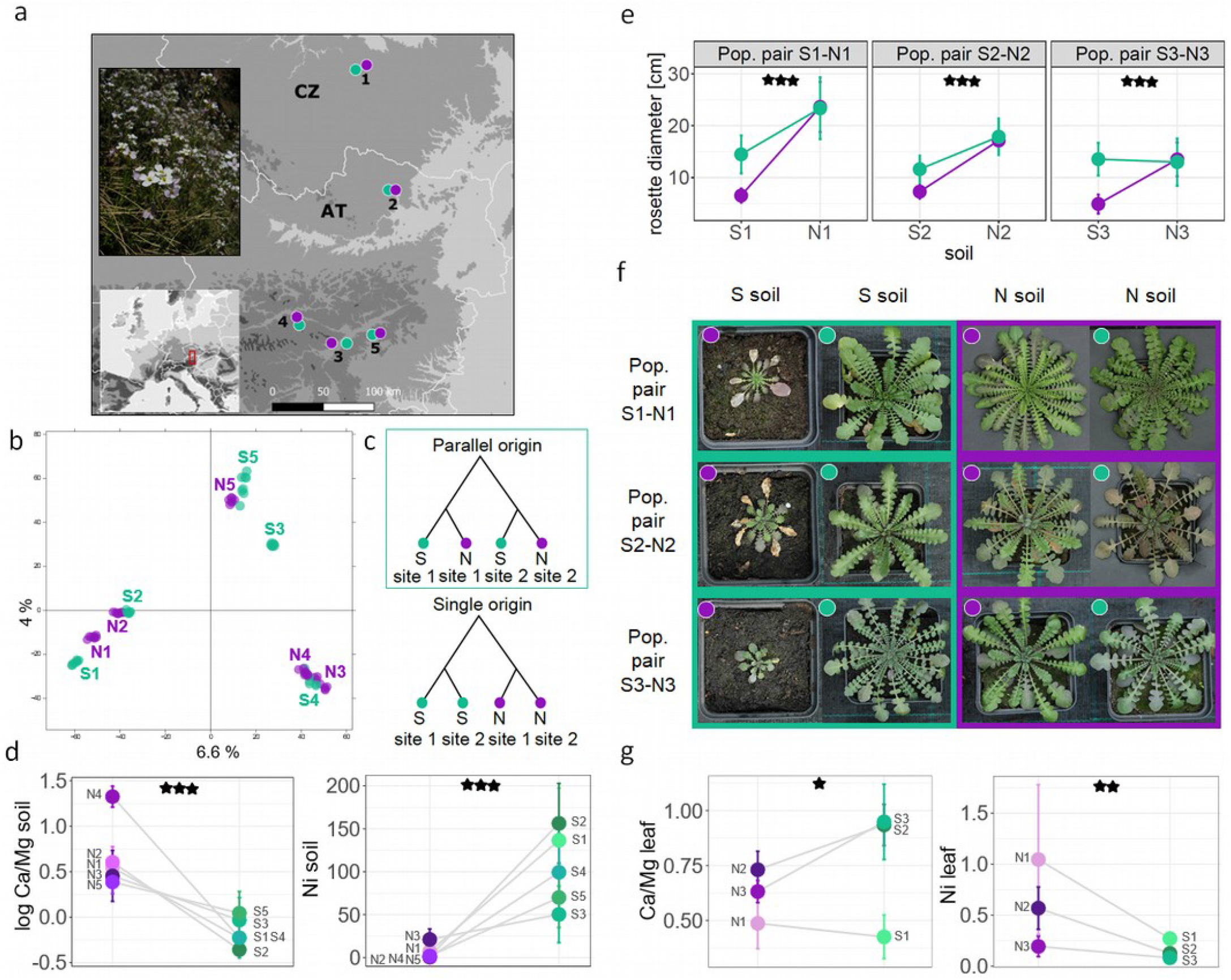
Parallel adaptation to challenging serpentine soils. **a)** Locations of the investigated serpentine (S, green) and non-serpentine (N, violet) populations sampled as spatially proximate pairs (numbers) in Central Europe with illustrative photo of an S population. **b)** Principal component analysis based on ∼1M fourfold-degenerate SNPs showing relationships among all individuals. **c)** Two contrasting evolutionary scenarios of serpentine colonization compared in coalescent simulations: in green frame is the topology receiving the highest support, consistently across all pairwise combinations of S-N population pairs. **d)** Differences in Ca/Mg ratio and in Ni concentrations [µg.g^-1^] in S and N soils from the original sampling sites (significant differences indicated as: F_1,77_ = 26.5, p < 0.001 and F_1,77_ = 117.4, p < 0.001 for Ca/Mg and Ni, respectively). **e)** Differences in maximum rosette size of three population pairs attained after three months of cultivation in local serpentine and non-serpentine substrates (significance of the soil treatment * soil origin interaction is indicated: F_1,90_ = 21.6, p < 0.001, F_1,96_ = 12.3, p < 0.001, and F_1,85_ = 42, p < 0.001 for population pairs 1, 2 and 3, respectively). **f)** Example photos illustrating parallel growth response in the three population pairs to serpentine soils (green frame) depending on the soil of origin (dot colour). **g**) Differences in ion uptake between originally S and N individuals when cultivated in serpentine soils; Ni concentrations were standardized by corresponding soil Ni values (One-way ANOVAs: F_1,27_ = 6.2, p = 0.019 and F_1,27_ = 13.5, p = 0.001 for Ca/Mg ratio and Ni, respectively). Significance: *p < 0.05, **p < 0.01, ***p < 0.001.

To assess whether the colonization of serpentines was accompanied by substrate adaptation we combined ionomics with a reciprocal transplant experiment. First, using ionomic profiling of native soil associated with each sequenced individual, we characterized major chemical parameters differentiating on both substrates (Fig 1d, Fig S3a, Fig S4). Among the 20 elements investigated (Fig S3a), only the bioavailable concentration of Mg, Ni, Co, and Ca/Mg ratio consistently differentiated both soil types (Bonferroni corrected one-way ANOVA taking population pair as a random variable). Serpentine sites were not macronutrient poor (Table S4) and were not differentiated from non-serpentines by bioclimatic parameters (annual temperature, precipitation, and elevation, Fig S3b), indicating that skewed Ca/Mg ratios and elevated heavy metal content are likely the primary selective agents on the sampled serpentine sites^35–37^.

We then tested for differential fitness response towards serpentine soil between populations of S vs. N origin using reciprocal transplant experiments. We cultivated plants from three population pairs (S1-N1, S2-N2, and S3-N3) on both native soil types within each pair for three months (until attaining maximum rosette size), observing significantly better germination and growth of the S plants in their native serpentine substrate as compared to their closest N relatives. First, we found a significant interaction between soil type and soil of origin at germination (GLM with binomial errors taking population pair as a random variable, Χ^2^ = 22.436, p < 0.001), although the fitness disadvantage of N plants in serpentine soil varied across population pairs (Fig S5). During subsequent cultivation, we recorded zero mortality but found a significant interaction effect between soil treatment and soil of origin on growth, as approximated by maximum rosette sizes (two-way ANOVA taking population pair as a random variable, F_1,277_ = 55.5, p < 0.001, Fig 1e; see Fig S6 for rosette size temporal development). Once again, the S plants consistently produced significantly larger rosettes (by 47% on average) than their N counterparts when grown in serpentine soil, indicating consistent substrate adaptation (Fig 1e, f). Finally, we evaluated differences in Ni and Ca/Mg accumulation in leaves harvested on plants cultivated in serpentine soils. Consistent with adaptive responses to soil chemistry, we found higher Ca/Mg ratio and reduced uptake of Ni (lower leaf/soil ratio) in tissue of serpentine plants relative to their non-serpentine counterparts (Fig 1g). Taken together, our demographic analysis complemented by transplant experiments support recent parallel serpentine adaptation of autotetraploid *A. arenosa* at five distinct sites, exhaustively covering all known serpentine populations of the species.

### Parallel genomic footprints of selection on serpentine at SNPs and TEs

Having determined that these five contrasts represent independent natural replicates of serpentine adaptation, we sought the genomic basis of these parallel adaptations. To do this, we first scanned for loci repeatedly exhibiting signals of selection upon each serpentine adaptation event in the complete resequencing dataset. We identified initial inclusive lists of gene coding loci (hereafter, ‘differentiation candidates’) exhibiting excessive differentiation between paired populations using 1% outlier F_ST_ window-based scans (490 to 525 candidate genes per pair; details in Methods; Dataset S2). Gene ontology (GO) enrichment of candidates from all five population pairs shows significant enrichment (FDR < 0.05) of functions considered relevant to serpentine adaptation^26,35^ such as cation transport, chemical homeostasis, post-embryonic development and transmembrane signalling receptor activity. Additionally, functional categories of proteins localized to the plasma membrane were enriched (Fig 2b; Dataset S3a).

**Fig 2.**
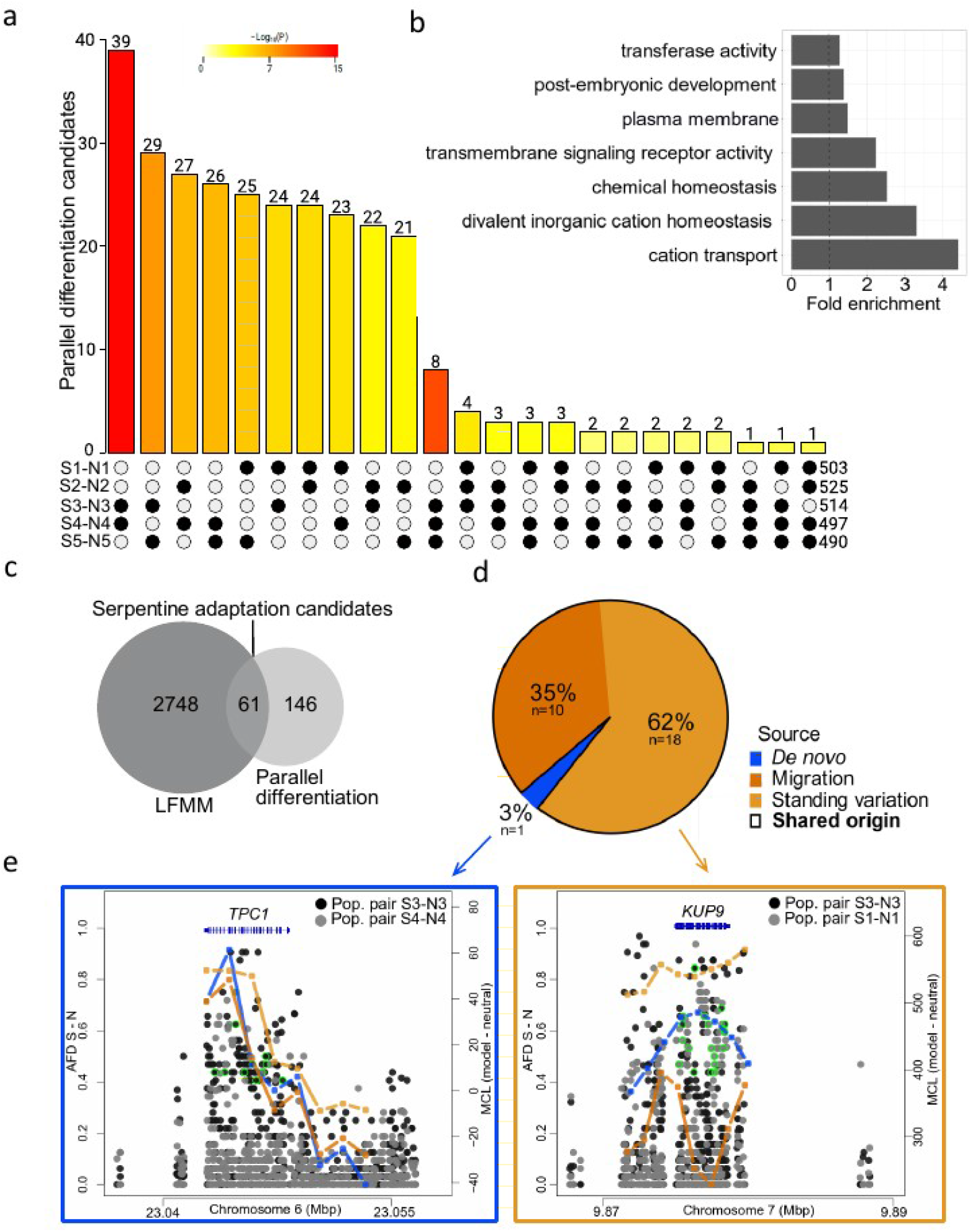
Parallelism in serpentine adaptation and sources of parallel adaptive variants. **a)** Intersection of differentiation candidates from each population pair (S1-N1 to S5-N5). All intersections were significant (p < 0.01, Fisher’s exact test, Dataset S3b). **b)** Gene ontology enrichment of the differentiation candidates (across all population pairs); for complete list of GO terms see Dataset S3a. **c)** Overlap between parallel differentiation candidates and Latent Factor Mixed Model (LFMM) candidates resulting in 61 serpentine adaptation candidate genes. **d)** Proportions of serpentine adaptation candidates originating from de novo mutations or being of shared origin out of the total of 29 cases of non-neutral parallelism as inferred by the Distinguishing among Modes of Convergence approach (DMC; see text for details). **e)** Two examples of parallel candidate loci, illustrating SNP divergence and maximum composite log-likelihood (MCL) estimation of the source of the selected alleles in DMC. Allele frequency difference (AFD) for locus with independent de novo mutations (left) and with parallel recruitment of shared ancestral standing variation (right). Left y-axis: AFD between S and N populations. Dots: AFD values of individual SNPs; bright green circles: non-synonymous SNPs with AFD ≥ 0.4; lines (right y-axis): maximum composite likelihood (MCL) difference between neutral vs. parallel selection scenario following colour scheme in panel 2d; gene models are in blue.

To greatly refine this broad list and pinpoint parallel evolution candidate genes, we overlapped the differentiation candidate gene lists across population pairs, identifying 207 ‘parallel differentiation candidates’ which represent divergence outliers in at least two S-N population pairs. The level of parallelism was greater than expected by chance (Fisher’s exact test; Fig 2a, Dataset S3b). The parallel differentiation candidates were significantly enriched (FDR < 0.05) for two GO terms: transporter activity (including ion transmembrane transporter and gated channel activity) and hydrolase activity (Dataset S3c).

As a complementary approach to the divergence scans, we identified alleles that consistently exhibited significant positive associations in allele frequency with key serpentine chemistry characteristics identified by ionomics (Ca/Mg ratio, high Mg, Ni, and Co) across every individual using latent factor mixed models (‘LFMM candidates’). We identified 2,809 genes harbouring ≥ 1 SNP significantly associated with at least one serpentine soil parameter (FDR < 0.05; Dataset S4). Finally, we overlapped these LFMM candidates with the parallel differentiation candidates to produce a final, most refined list of 61 ‘serpentine adaptation candidates’ (Fig 2c, Fig S7, Dataset S5, Dataset S6a). This conservative approach aims to identify the strongest candidates underlying serpentine adaptation. Although it discards population-specific (private) candidates and cases of distinct genetic architecture of a trait (e.g. distinct genes affecting same pathway) it also importantly minimizes false positives from population-specific selection and genetic drift.

These 61 serpentine adaptation candidates were significantly enriched (FDR < 0.05) for categories related to ion and transmembrane transport, and specifically, voltage-gated calcium channel activity (Dataset S6b). Candidates included the *NRT2*.*1* and *NRT2*.*2* high-affinity nitrate transporters, which act as a repressors of lateral root initiation^39,40^; *RHF1A*, which is involved in gametogenesis and transferase activities^41,42^; *TPC1*, a central calcium channel which mediates plant-wide stress signalling and tolerance^43^; and potassium transporters *AKT5* and *KUP9*. Furthermore, when we compared our serpentine adaptation candidates (n=61) to candidate loci for parallel serpentine adaptation in *A. lyrata* (n=62) from a previous study^44^ we found only two loci in common (significant overlap; p < 0.007), *KUP9* and *TPC1*, further supporting important roles of these two ion transporters in repeated adaptation to serpentine soil.

Single nucleotide polymorphism data only present part of the picture, and despite linkage, do not directly capture changes mediated by structural variation. Thus, we also investigated divergence at transposable elements (TE) variations in population pairs 1 to 4 (relatively lower coverage of the N5 population did not permit this analysis) based on 21,690 TE variants called using the TEPID approach that is specifically designed for population TE variation studies^45^ (Dataset S7). Assuming linkage between TE variant and surrounding SNPs, we then applied a similar differentiation outlier window-based workflow as specified above and identified 92–115 TE-associated differentiation candidate genes per S-N contrasts (Dataset S8). We found in total 23 enriched gene categories (Dataset S9; sum of GO terms across population pairs) that included genes involved in transport and water transporter activity. By overlapping these candidate lists across S-N pairs, we identified 13 parallel TE-associated differentiation candidates (Fig S8, Dataset S8f; significant overlap, p < 0.05). These loci included the plasma membrane protein *PIP2*, the putative apoplastic peroxidase *PRX37*, and *RALF-LIKE 28*, which is involved in calcium signalling. This suggests a potential impact of TEs on serpentine adaptation and gives discrete candidates for future study.

### Sources of adaptive variation

Next, we tested whether variants in each serpentine adaptation candidate have arisen by parallel *de novo* mutations or instead came from pre-existing variation shared across populations. To do so, we modelled allele frequency covariance around repeatedly selected sites for each locus and identified the most likely of the four possible evolutionary scenarios using a designated ‘Distinguishing among Modes of Convergence’ (DMC) approach^38^: (i) a null-model assuming no selection (neutral model) (ii) independent *de novo* mutations at the same locus; (iii) repeated sampling of shared ancestral variation; and (iv) sharing of adaptive variants via migration between adapted populations. For simplicity, we consider scenarios (iii) and (iv) jointly as ‘shared origin’ because both processes operate on alleles of a single mutational origin, in contrast to scenario (ii). To choose the best fitting scenario for each of the 61 candidate genes, we compared the maximum composite log-likelihoods (MCL) between the four scenarios (see Methods; Table S5). This analysis indicated that parallel selection exceeded the neutral model for 29 out of the total 84 candidate cases of parallelism (i.e. cases when two population pairs shared one of the 61 serpentine adaptation candidates). Shared origin dominated our non-neutral models of adaptive variation, representing 97% of the non-neutral cases (28/29 cases; Fig 2d and Dataset 6a). The alternate non-neutral scenario, parallel *de novo* origin, was supported only for a single locus, TWO PORE CHANNEL 1 (*TPC1*) in one case (S3-N3 and S4-N4 pairs). Using a more permissive threshold for identifying differentiation candidates (3% outliers, leading to a five-fold increase in parallel candidates) resulted in a similar DMC estimate of the proportion of the shared variation scenario (103/114 cases, i.e. 90%; Dataset S10, Fig S9), indicating that our inference of the dominant role of shared variation is not dependent on a particular outlier threshold. Finally, we applied similar approach to parallel TE-associated differentiation candidates (n=13) assuming selection on TE variants left footprint in surrounding SNP-allele frequency covariances. We found a single non-neutral candidate, *ATPUX7*, for which parallel selection on standing variation was inferred (Dataset S8f). In summary, by a combination of genome-wide scanning with a designated modelling approach, we find that a non-random fraction of loci is likely re-used by selection on serpentine, sourcing almost exclusively from a pool of alleles shared across the variable autotetraploid populations.

### Rapid recruitment of convergent de novo *mutations at the calcium channel TPC1*

One advantage of the DMC approach is an objective model selection procedure. However, it does not give fine scale information about the distribution of sequence variation at particular alleles. Therefore, we further investigated candidate alleles of the *TPC1* gene, for which DMC results suggested the sweep of different *de novo* mutations in independent serpentine populations. Upon closer inspection of all short-read sequences complemented by Sanger sequencing of additional 40 individuals from the three serpentine populations, a remarkably specific selection signal emerged. We found two absolutely serpentine-specific, high-frequency, non-synonymous mutations only at residue 630, overlapping the region of the highest MCL estimate for the *de novo* scenario in DMC, and directly adjacent to the selectivity gate of the protein in structural homology models (Fig 3; Table S6). Of the two, the polymorphism Val630Leu is nearly fixed in the S3 population (25 homozygous Leu630 individuals and four heterozygous Leu630/Val630 individuals out of 29 individuals), and is at a high frequency in the S5 population (one homozygous Leu630 individual and 17 heterozygous out of 20 individuals); the second convergent Val630Tyr mutation is at high frequency in the S4 population (four homozygous Tyr630 individuals and 18 heterozygous Tyr630/Val630 out of 25 individuals; Fig 3a, b and Fig S10). Strikingly, Val630Tyr requires a three-nucleotide mutation covering the entire codon (GTA to TAT). Neither of these variants were found in any other *A. arenosa* population in a range-wide catalogue^8^ encompassing 1,724 *TPC1* alleles (including 368 alleles from the focal area of Eastern Alps; Fig 3a, Dataset S11) nor in available short read data of the other two *Arabidopsis* outcrossing species (224 *A. lyrata* and 178 *A. halleri* alleles, respectively, Fig S11), indicating that both are private to serpentine populations. Altogether, the absolute lack of either *A. arenosa* serpentine-specific variant in non-serpentine sampling across the genus strongly supports the conclusions of the DMC modelling of their independent *de novo* mutation origin.

**Fig 3.**
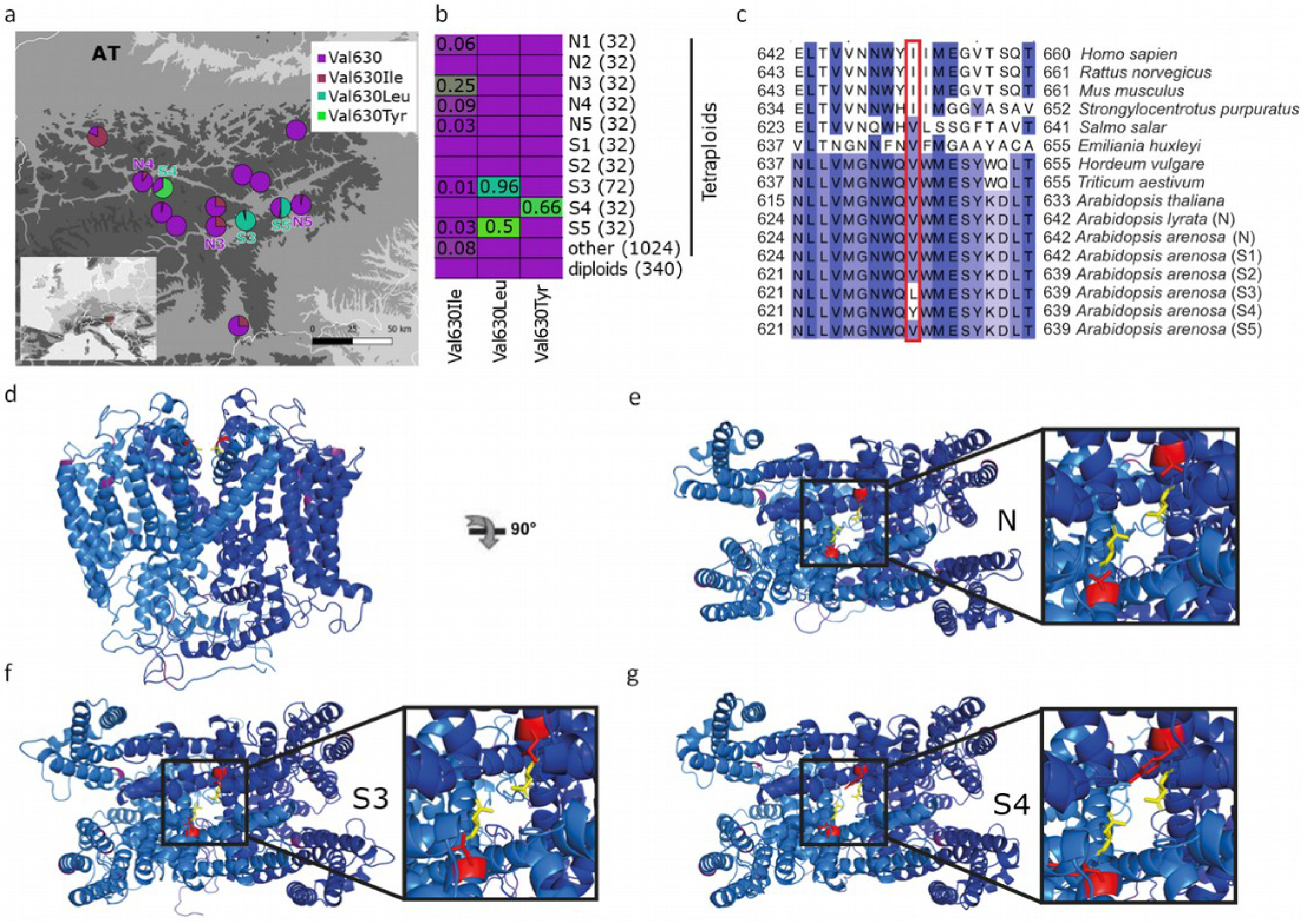
Serpentine-private, parallel de novo high impact protein changes in TPC1 locus. **a)** All populations with serpentine specific variants and all other resequenced A. arenosa populations in the focal area of Eastern Alps, showing frequencies of amino acid substitutions at residue 630 as pie charts (Dataset S11). **b)** Population frequencies of substitutions in the residue 630 among 1,724 alleles from range-wide A. arenosa resequenced sample. Colours denote frequencies from ancestral non-serpentine (violet) to serpentine-specific alleles (green) and number in brackets denotes total N of alleles screened (Dataset S11). **c)** Cross-kingdom conservation of the site shown by multiple sequence alignment of surrounding exon including consensus sequences (AF > 0.5) from all serpentine A. arenosa populations (in S5 population the frequency of Val630Leu is 0.5). Residues are coloured according to the percentage that matches the consensus sequence from 100% (dark blue) to 0% (white), the position of the serpentine-specific high frequency non-synonymous SNP is highlighted in red. **d-g)** Structural homology models of A. arenosa TPC1 alleles. Dimeric subunits are coloured blue or marine. Non-synonymous variation that is not linked to serpentine soil is coloured deep purple. Residue 630 is coloured red and drawn as sticks. The adjacent residue, 627 (631 in A. thaliana), which has an experimentally demonstrated key role in selectivity control, is yellow and drawn as sticks. **d)** Side view of the non-serpentine allele. **e)** Top view of the non-serpentine allele with the detail of the pore opening depicted in the inset. **f)** Top view of the Leu630 allele private for S3 and S5 populations. **g)** Top view of the Tyr630 allele private for S4 population.

To investigate the potential functional impact of these high frequency, convergent amino acid changes, we first generated an alignment for *TPC1* homologs across the plant and animal kingdoms. Residue 630 (634/633 in *A. thaliana/A. lyrata*) is conserved as either a Val or Ile across kingdoms, except for the serpentine *Arabidopsis* populations (Fig 3b, c, Fig S12). Val/Ile and Tyr are disparate amino acids in both size and chemical properties, so this substitution has high potential for a functional effect. Although the difference between Val/Ile and Leu is not as radical, we predicted that the physical difference between the side chains of Val/Ile versus Leu (the second terminal methyl group on the amino acid chain) may also have a functional effect by making new contacts in the tertiary structure. To test this, we performed structural homology modelling of all alleles found in *A. arenosa* (Fig 3d-g), using two crystallographically-determined structures as a template (PDB codes 5DQQ and 5E1J^46,47^). In the tertiary structure, residue 630 sits adjacent to the Asn residue (Asn627 in *A. arenosa*), which forms the pore’s constriction point and has been shown to control ion selectivity in *A. thaliana*^46–49^. In *A. thaliana* this Asn627 residue, when substituted by site-directed mutagenesis to the human homolog state can cause Na+ non-selective *A. thaliana TPC1* to adopt the Na+ selectivity of human *TPC1*^47^. Depending on the rotameric conformation adopted, the Leu630 allele forms contacts with the selectivity-determining Asn627 residue, while the non-serpentine Val630 does not (Fig 3e).

Our modelling suggests that the Tyr630 residue is even more disruptive: the Tyr side chain can adopt one of two broad conformations, either sticking into the channel, where it occludes the opening, or sticking away from the channel and directly into surrounding residues, which have been shown to be important for stabilising Asn627 in *A. thaliana*. Finally, to determine likely rotameric conformations, we generated 100 models of the S4 homodimer (two Tyr630 alleles). In this case, the two residues were significantly less likely to both point into the channel (Table S7), suggesting that this is difficult for the structure to accommodate. Because the S4 Tyr630 allele is predominantly in a heterozygous state with the Val630 allele in nature, we also generated 100 models of the S4 heterodimer (one Tyr630 and one Val630 allele). In the heterodimer there was no significant difference between the occurrence of the two side-chain conformations (Table S7), suggesting that either disruptive conformation, sticking into the pore or affecting nearby functional residues, is possible. Taken together, these results suggest that even the presence of a single Tyr630 variant within a dimer will have a substantial impact on *TPC1* function, and that the Tyr630 allele is likely to be dominant or partially-dominant. This prediction is consistent with its predominant occurrence in a heterozygous state and a ∼50% allele frequency in S4 (Fig S10). We also speculate that if homodimers of the Tyr allele are too disruptive to *TPC1* function this may result in a heterozygote advantage maintained by balancing selection. In conclusion, our results suggest that the serpentine-linked allelic variation at residue 630 impacts the selectivity of *TPC1*, with functional implications, and that independent convergent *de novo* mutations have been repeatedly selected upon during adaptation to serpentine soils.

## Discussion

Here we inferred the evolutionary sources of adaptive variation in autotetraploid populations by taking advantage of naturally replicated adaptation to toxic serpentine soil in wild *Arabidopsis arenosa*. Using a designated statistical approach leveraging parallelism, we inferred that nearly all parallel adaptation candidates were sourced from a common pool of alleles that was shared across populations. However, for one exceptional candidate — a central calcium channel shown to mediate stress signalling *TPC1*^43^ — we identified independent *de novo* mutations at the same otherwise highly conserved site with likely functional consequences.

The potential of polyploidy to enhance adaptation is a matter of ongoing debate that is mainly fuelled by theory-based controversies revolving around the efficiency of selection^6,19,50,51^ and observations of the frequencies of WGD events in space and time^5,22^. In contrast, empirical population-level investigations that unravel evolutionary mechanisms operating in natural autopolyploids are very scarce. It has been shown that autopolyploids may rapidly react to new challenges by landscape genetic^23^ and experimental studies^24,52^. Our transplant experiments coupled with demographic investigations support this and further demonstrate that such rapid adaptation may be repeated many times within a species, drawing on the same variants. Furthermore, a previous study in *A. arenosa* showed that the genome-wide proportion of non-synonymous polymorphisms fixed by directional selection was higher in tetraploids than diploids, suggesting increased adaptive variation in natural autopolyploids^8^. However, it remains unclear if such variation reflects increased input of novel mutations (as observed, e.g. in experimental yeast populations^17^), or sampling from increased standing variation (as predicted by theory^5,6^). Here we find nearly exclusive sampling from a shared pool of variants. Such a dominant role of pre-existing variation is in line with the studies of parallel adaptation in diploid systems such as *Littorina* snails^53,54^, stickleback^55,56^, and *Helliconius* butterflies^57^. Yet examples are lacking from autopolyploid systems, where larger pool of standing variation is expected due to larger effective population size and polysomic masking of allelic variation^5,58^. Sharing alleles that have persisted in a specific genomic environment already for some time may be particularly beneficial under an intense selection when rapid and efficient adaptive response is needed^59–62^. In addition, standing genetic variation likely minimizes negative pleiotropic effects of linked adaptive variants^62–64^. It should be noted, however, that our estimates may be biased upward for shared alleles by focusing only on cases of parallelism, which provided testable framework for our inference of the evolutionary sources. Larger fractions of novel mutations may be represented among the non-parallel adaptive variation, which is, however, harder to identify.

In contrast to shared variants, empirical evidence for parallel *de novo* mutations within species is rare even in diploids^65–67^ and we lack any example from polyploids. Theory suggests that such a scenario is unlikely for autopolyploids, as reduced efficacy of selection on a novel, initially low-frequency variants is predicted for most dominance states in autopolyploids^6,18,19^. On the other hand, beneficial alleles are introduced at increased rates in doubled genomes^6,17^ and additional variation may accumulate due to polysomic masking^6^. Here we provide an example of parallel recruitment of two distinct *de novo* mutations with likely phenotypic effect in separate polyploid populations within a species, demonstrating that adaptive sourcing from novel polymorphisms is in fact feasible in even in autopolyploids. Interestingly, high frequencies of homozygous individuals in one population demonstrates that such novel variants may approach fixation, in stark contrast to theory, which expect incomplete sweeps of dominant mutations to be prevalent in autotetraploids^6,19^. On the other hand, the prevalence of heterozygotes in the other serpentine population together with results of structural modelling suggest (at least partial) dominance of the serpentine allele.

Overall, our study demonstrates that rapid environmental adaptation may repeatedly occur in established autopolyploid populations, dominantly sourcing from a large pool of pre-existing variation, yet exceptionally also from recurrent *de novo* mutations. Thus, these results support the emerging view of autopolyploids as diverse evolutionary amalgamates, capable of flexible adaptation in response to environmental challenge.

## Methods

### Field sampling

Serpentines occur in Central Europe as scattered edaphic ‘islands’ surrounded by open rocky habitats on other substrates in which autotetraploid *A. arenosa* frequently occur. In contrast, *A. arenosa* colonised only some serpentine sites in this area^68,69^ indirectly suggesting that colonisation of serpentine sites by surrounding non-serpentine populations happened in parallel and was probably linked with local substrate adaptation. To test this hypothesis, we sampled all five serpentine (S) populations of *A. arenosa* known to date and complemented each by a proximal (< 19 km distant) non-serpentine (N) population. All N populations grew in similar vegetation (rocky outcrops in open forests or grasslands) and soil type (siliceous to neutral rocks; Table S1). Diploid serpentine *A. arenosa* is not known, even though serpentine barrens are frequent in some diploid-dominated areas such as the Balkan peninsula. The sampled populations covered considerable elevational gradient (414-1750 m a.s.l.), but the differences in elevation within the pairs were small except for one pair where no nearby subalpine non-serpentine population exists (population pair S4-N4, difference 740 m).

We sampled eight individuals per every population for genomic analysis and confirmed their tetraploid level by flow cytometry. For each individual, we also sampled soil from very close proximity to the roots (∼10 to 20 cm below ground), except for N5 population for which genotyped data were already taken from the previous study^8^. There, we collected an additional eight soil samples and use their average in following environmental association analysis. For the transplant experiment, we also collected seeds (∼20-30 maternal plants/population) from three population pairs (S1-N1, S2-N2, S3-N3) and bulks of soil ∼80 l (sieved afterwards) from the natural sites occupied by these six populations.

### DNA extraction, library preparation, sequencing, raw data processing and filtration

We stored all samples for this study in RNAlater (R0901-500ML, SIGMA-ALDRICH CO LTD) to avoid genomic DNA degradation and we further prepared the leaf material as described in^70–72^. We extracted DNA as described in Protocols. Genomic libraries for sequencing were prepared using the Illumina TRUSeq PCR-free library. Libraries were sequenced as 150 bp paired end reads on a HiSeq 4000 (3 lanes in total) by Norwegian Sequencing Centre, University of Oslo.

We used trimmomatic-0.36^73^ to remove adaptor sequences and low-quality base pairs (< 15 PHRED quality score). Trimmed reads longer than 100 bp were mapped to reference genome of North American *Arabidopsis lyrata*^74^ by bwa-0.7.15^75^ with default setting. Duplicated reads were identified by picard-2.8 (https://github.com/broadinstitute/picard) and discarded together with reads that showed low mapping quality (< 25). Afterwards we used GATK v.3.7 to call and filter reliable variants and invariant sites according to best practices^76^ (complete variant calling pipeline available at https://github.com/vlkofly/Fastq-to-vcf). Namely, we used the HaplotypeCaller module to call variants per individual using the ploidy = 4 option which enables calling full tetraploid genotypes. Then, we aggregated variants across all individuals by module GenotypeGVCFs. We selected only biallelic SNPs and removed those that matched following criteria: Quality by Depth (QD) < 2.0, FisherStrand (FS) > 60.0, RMSMappingQuality (MQ) < 40.0, MappingQualityRankSumTest (MQRS) < −12.5, ReadPosRankSum < −8.0, StrandOddsRatio (SOR) > 3.0. We called invariant sites also with the GATK pipeline similarly to variants, and we removed sites where QUAL was lower than 15. Both variants and invariants were masked for sites with average read depth higher than 2 times standard deviation as these sites were most likely located in duplicated regions and we also masked regions with excessive heterozygosity, representing likely paralogous mis-assembled regions, following ref. ^8^. One individual per each S2 and N5 populations was excluded due to exceptionally bad data quality (low percentage of mapped reads and low read depth, <10 on average), leaving us with a final dataset of 78 individuals that were used in genomic analyses. This pre-filtered dataset contained 110,358,565 sites (of which 11,744,200 were SNPs) with average depth of coverage 21x (Dataset S1a).

### Reconstruction of population genetic structure

We inferred the population genetic structure, diversity and relationships among individuals from putatively neutral fourfold-degenerate (4dg) SNPs filtered for read depth (DP) > 8 per individual and maximum fraction of filtered genotypes (MFFG) of 0.2, i.e. allowing max. 20% missing calls per site (1,042,793 SNPs with a total of 0.49 % missing data; see Table S1, S2, and S3 for description of datasets and filtration criteria). We used several complementary approaches. First, we ran principal component analysis (PCA) on individual genotypes using glPCA function in adegenet v.2.1.1 replacing the missing values by average allele frequency for that locus. Second, we applied model-based clustering with accelerated variational inference in fastStructure v.1.0^77^. To remove the effect of linkage, we randomly selected one SNP per a 1 kb window, keeping 10kb distance between the windows and, additionally, filtered for minimum minor allele frequency (MAF) = 0.05 resulting in a dataset of 9,923 SNPs. As fastStructure does not handle the polyploid genotypes directly, we randomly subsampled two alleles per each tetraploid site using a custom script. This approach has been demonstrated to provide unbiased clustering in autotetraploid samples in general^78^ and *Arabidopsis* in particular^8^. We ran fastStructure with 10 replicates under K=5 (corresponding to the number of population pairs) with default settings. Third, we inferred relationships among populations using allele frequency covariance graphs implemented in TreeMix v.1.13^79^. We used custom python3 scripts (available at https://github.com/mbohutinska/TreeMix_input) to create the input files. We ran TreeMix analysis rooted with an outgroup population (tetraploid *A. arenosa* population ‘Hranovnica’ from the area of origin of the autotetraploid cytotype in Western Carpathians^34^, Dataset S1a for samples overview). We repeated the analysis over the range of 0–6 migration edges to investigate the optimal number of informative migration events (Fig S1c). We bootstrapped the scenario without migration (the topology did not change with adding the migrations) choosing bootstrap block size 1 kb (the same window size also for the divergence scan, see below), 100 replicates and summarized the results using *SumTrees*.*py* function in DendroPy^79^. Finally, we calculated nucleotide diversity (π) and Tajima’s D for each) and Tajima’s D for each population and pairwise differentiation (F_ST_) for each population pair using custom python3 scripts (available at https://github.com/mbohutinska/ScanTools_ProtEvol; see Table S3 for number of sites per each population). For nucleotide diversity calculation we down-sampled each population to six individuals on a per-site basis to keep equal sample per each population while also keeping the maximum number of sites with zero missingness.

### Demographic inference

We performed demographic analyses in fastsimcoal v.2.6^80^ to specifically test for parallel origin of serpentine populations and to estimate divergence time between serpentine and proximal non-serpentine populations. We constructed unfolded multidimensional site frequency spectra (SFS) from the variant and invariant 4dg sites (filtered in the same ways as above, Table S2) using custom python scripts published in our earlier study (FSC2input.py at https://github.com/pmonnahan/ScanTools/)^8^. We repolarized a subset of sites using genotyped individuals across closely related diploid *Arabidopsis* species to avoid erroneous inference of ancestral state based on a single reference *A. lyrata* individual following ref. ^8^.

First, we tested for parallel origin of serpentine populations using population quartets (two pairs of geographically proximal serpentine and non-serpentine populations) and iterated such pairs across all combinations of regions (10 pairwise combinations among the five regions in total). For each quartet we created four-dimensional SFS and compared following four evolutionary scenarios (Fig. S2): (i) parallel origin of serpentine ecotype – sister position of serpentine and non-serpentine populations within the same region, (ii) parallel origin with migration – the same topology with additional gene flow between serpentine and the proximal non-serpentine population, (iii) single origin of serpentine ecotype – sister position of serpentine populations and of non-serpentine populations, respectively, (iv) single origin with migration – the same topology with additional gene flow between serpentine and the proximal non-serpentine populations. For each scenario and population quartet, 50 fastsimcoal runs were performed. For each run, we allowed for 40 ECM optimisation cycles to estimate the parameters and 100,000 simulations in each step to estimate the expected SFS. We used wide range of initial parameters (effective population size, divergence times, migration rates; see the example *.est and *.tpl files provided for each model tested in the Dataset S12) and assumed mutation rate of 4.3×10^−8^ inferred for *A. arenosa* previously^32^. Further, we extracted the best likelihood partition for each fastsimcoal run, calculated Akaike information criterion (AIC) and summarized the AIC values across the 50 fastsimcoal runs. The scenario with lowest median AIC values within each particular population quartet was preferred (Fig S2).

Second, we estimated divergence time between S-N populations from each population pair (i.e. S1-N1, S2-N2, S3-N3, S4-N4, and S5-N5) based on two-dimensional SFS using the same fastsimcoal settings as above. We simulated according to models of two-population split, not assuming migration because the model with migration did not significantly increase the model fit across the quartets of populations (Fig S2). To calculate 95% confidence intervals for parameter estimates (Table 1) we sampled with replacement the original SNP matrices to create 100 bootstrap replicates of the two-dimensional SFS per each of the five population pairs.

### Window-based scans for directional selection

We leveraged the fivefold-replicated natural setup to identify candidate genes that show repeated footprints of selection across multiple events of serpentine colonization. First, we identified genes of excessive divergence for each pair of proximal serpentine (S) – non-serpentine (N) populations (five pairs in total). We calculated pairwise F_ST_^81^ for non-overlapping 1 kb windows along the genome with the minimum of 10 SNPs per window using the custom script (https://github.com/mbohutinska/ScanTools_ProtEvol) based on ScanTools pipeline that has been successfully applied in our previous analyses of autotetraploid *A. arenosa*^8^. The window size is larger than average genome-wide LD decay of genotypic correlations (150-800 bp) estimated in *A. arenosa*^82^. We identified the upper 99% quantile of all 1 kb windows in the empirical distribution of F_ST_ metric per each population pair. Then, we identified candidate genes for directional selection (‘differentiation candidates’) as genes overlapping with these 1% outlier windows using *A. lyrata* gene annotation^83^, where the gene includes 5’ UTRs, start codons, exons, introns, stop codons, and 3’ UTRs. In total, we identified 2,245 such genes in all five population pairs. Finally, we greatly refined this inclusive list and identified parallel differentiation candidates as overlapping genes in differentiation candidate lists across at least two population pairs. We tested if such overlap is higher than a random number of overlapping items given the sample size using Fisher’s exact test in SuperExactTest R package^84^. Our F_ST_ based detection of outlier windows was not largely biased towards regions with low recombination rate (based on the available *A. lyrata* recombination map^85^; Fig S13).

### Environmental association analysis

As a complementary approach for detection of serpentine selection candidates we performed environmental association analysis using latent factor mixed models - LFMM 2 (https://bcm-uga.github.io/lfmm/)^8 6^. We tested the association of allele frequencies at each SNP with associated soil concentration of the key elements differentiating serpentine and non-serpentine soils: Ca/Mg ratio, and bioavailable soil concentrations of Co, Mg, and Ni. Only those elements were significantin one-way ANOVAs (Bonferroni corrected) testing differences in elemental soil concentration between S and N population, taking population pair as a random variable. We retained 1,783,055 SNPs without missing data and MAF > 0.05 as an input for the LFMM analysis. LFMM accounts for discrete number of ancestral population groups as latent factors - we used five latent factors corresponding to five population pairs in our dataset. To identify SNPs significantly associated with soil variables, we transformed p-values to false discovery rate (FDR < 0.05) based on q-values using the qvalue R package v.2.20^87^. Finally, we annotated the candidate SNPs to genes, termed ‘LFMM candidates’ (at least one significantly associated SNP per candidate gene).

We made a final shortlist of serpentine adaptation candidates by overlapping the LFMM candidates, reflecting significant association with important soil elements, with the previously identified parallel differentiation candidates, mirroring regions of excessive differentiation repeatedly found across parallel population pairs. For visualisation purposes (Fig 2e), we annotated SNPs in the serpentine adaptation candidates using SnpEff 4.3^88^ following *A. lyrata* version 2 genome annotation^83^.

### Transposable element (TE) variant calling and analysis

TE variants (insertions or deletions) among sequenced individuals were identified and genotyped in population pairs 1-4 using TEPID v.0.8^45^ following the approach described in ref. ^89^ (relatively lower coverage of the N5 population did not permit this analysis in the last pair). We annotated TEs based on available *A. lyrata* TE reference^90^. TEPID is based on split and discordant read mapping information and employs read mapping quality, sequencing breakpoints and local variation in sequencing coverage to call absence of reference TEs as well as the presence of non-reference TE copies. This method is specifically suited for studies at the population level as it takes intra-population polymorphism into account to refine TE calls in focal samples by supporting reliable call of non-reference alleles under lower thresholds when found in other individuals of the population. We filtered the dataset by excluding variants with MFFG > 0.2 and DP < 8, which resulted in 21,690 TE variants (13,542 deletions and 8,148 insertions as compared to the reference).

Assuming linkage between TE variants and nearby SNPs^44^, we calculated pairwise F_ST_^81^ using SNP frequencies (in the same way as specified above) for non-overlapping 1 kb windows containing TE variant(s) for each population pair. The candidate windows for directional selection were identified as the upper 99% quantile of all windows containing a TE variant in the empirical distribution of F_ST_ metric per each population pair. Further, we identified candidate genes (‘TE-associated differentiation candidates’) as those present up to +-2kb upstream and downstream from the candidate TE variant (assuming functional impact of TE variant until such distance, following ref. ^92^). Finally, we identified parallel TE-associated differentiation candidates as those loci that appeared as candidates in at least two population pairs.

### Gene Ontology (GO) enrichment analysis

We inferred potential functional consequences of the candidate gene lists using gene ontology (GO) enrichment tests using BiNGO v.3.0.3^93^ in CYTOSCAPE version 3.7.1 with the GO information associated with orthologous *A. thaliana* gene identifiers. We examined GO terms involved in molecular functions, biological processes, and cellular components. We considered GO terms significantly enriched if FDR p-adjusted values (Benjamini-Hochberg correction) were ≤ 0.05 for every SNP data analysis and < 0.1 for TE-associated data analysis in order to compensate for lower number (∼5-times less) of candidate genes in the latter dataset. We summarized the results and removed redundant enriched GO terms using REVIGO^88^.

### Modelling the sources of adaptive variation

For each serpentine adaptation candidate (n=61 and n=13 identified using SNPs and TE-associated variants, respectively), we modelled whether it exhibits patterns of parallel selection that is beyond neutrality and if so whether the parallel selection operated on *de novo* mutations or rather called on pre-existing variation shared across populations. We used model-based likelihood approach that is specifically designed to identify loci involved in parallel evolution and to distinguish among their evolutionary sources (Distinguishing Among Modes of Convergence; DMC^38^). Convergent is analogous to parallel in this case as the entire approach is designed for closely related populations.

We considered the following four evolutionary scenarios assuming distinct variation sources: (i) no selection (neutral model), (ii) independent *de novo* mutations at the same locus, (iii) repeated sampling of ancestral variation that was standing in the non-adapted populations prior the onset of selection, and (iv) transfer of adaptive variants via gene flow (migration) from another adapted population. We interpreted the last two scenarios together as a variation that is shared across populations as both operate on the same allele(s) that do not reflect independent mutations.

We estimated composite log-likelihoods (CLs) for each gene and under each scenario using a broad range of realistic parameters taking into account demographic history of our populations inferred previously and following recommendations in ref. ^38^ (positions of selected sites, selection coefficients, migration rates, times for which allele was standing in the populations prior to onset of selection, and initial allele frequencies prior to selection; see Table S5 for the summary of all parameters and their ranges). We placed the positions of selected sites at eight locations at equal distance along each serpentine adaptation candidate gene. For the calculation of co-ancestry decay, we also considered a 25 kb upstream and downstream region from each gene. To choose the best fitting scenario for each candidate, we compared the maximum composite log-likelihoods (MCL) between the parallel selection scenario and neutral scenario. We considered the MCL difference between a parallel and neutral model significant if it was higher than the maximum of the distribution of the differences from the simulated neutral data in *A. arenosa* inferred in ref. ^82^ (i.e. 21, a conservative estimate). Further, to focus only on divergence caused by selection in serpentine populations (i.e. eliminating divergence signals caused by selection in N populations), only cases of selection in the serpentine populations in which the scenario of parallel selection had a considerably higher MCL estimate (>10%) than selection in non-serpentine populations were taken into account.

To ensure that our inference on the relative importance of shared vs. *de novo* variation is not biased by arbitrary outlier threshold selection, we re-analyzed the SNP dataset using a 3% outlier F_ST_ threshold for identifying differentiation candidates. We overlapped the resulting 1179 parallel differentiation candidates with the LFMM candidates and subjected the resulting 246 serpentine adaptation candidates to DMC modelling in the same way as described above (Dataset S10). For each of the 246 genes and parallel quartets of populations (in total 420 cases of parallelism) we again compared the four scenarios as described above (Fig S9; Dataset S10).

### Reciprocal transplant experiment

To test for local substrate adaptation in three serpentine populations we compared plant fitness in the native vs. foreign soil in a reciprocal transplant experiment. We reciprocally transplanted plants of serpentine and non-serpentine origin from three population pairs (S1-N1, S2-N2, S3-N3) that served as representatives of independent colonization in each broader geographic region (Bohemian Massif, lower Austria and Eastern Alps, respectively). As the most stressful factor for *A. arenosa* populations growing on serpentine sites is the substrate (ref. ^29^ and Fig. S3a, b), we isolated the soil effect by cultivating plants in similar climatic conditions in the greenhouse.

For each pair we cultivated the plants in serpentine and non-serpentine soil originating from their original sites (i.e. S1 plant cultivated in S1 and N1 soil and vice versa) and tested for the interaction between the soil treatment and soil of origin in selected fitness indicators (germination and rosette diameter sizes). We germinated seeds from 12 maternal plants (each representing a seed family of a mixture of full- and half-sibs) from each population in petri dishes filled by either type of soil (15 seeds/family/treatment). Seeds germinated in the growth chamber (Conviron) under conditions approximating spring season at the original sites: 12 h dark at 10°C and 12 h light at 20°C. We recorded the germination date as the appearance of cotyledon leaves for the period of 20 days after which there were no new seedlings emerging. We tested for the effect of substrate of origin (serpentine versus non-serpentine), soil treatment (serpentine versus non-serpentine) and their interaction on germination proportion using GLM with binomial errors, taking population pair (1-3) as a random factor.

Due to zero germination of N1 seeds in S1 soil, we measured differential growth response on plants that were germinated in the non-serpentine soils and were subjected to the differential soil treatment later, in a seedling stage. We chose 44-50 seedlings equally representing progeny of 11 maternal plants per each population (in total 284 seedlings), transferred each plant to a separate pot filled either with ∼1 l of the original or the alternative paired soil (i.e., S1 soil for N1 population and vice versa). We randomly swapped the position of each pot twice a week and watered them with tap water when needed. We measured the rosette diameter and counted number of leaves (which correlated with the rosette diameter, R=0.85, p < 0.001) twice a week for five weeks until rosette growth reached a plateau (Fig S6). By that time, we observed zero mortality and only negligible flowering (1%, 4 of 284 plants). We tested if soil treatment (serpentine versus non-serpentine) with the interaction of soil of origin (serpentine versus non-serpentine) had a significant effect on rosette diameter sizes (the maximum rosette diameter sizes from the last the 10^th^ measurement as a dependent variable), using two-way ANOVA taking population pair (1-3) as a random factor.

### Elemental analysis of soil and leaf samples

We quantified the soil elemental composition by Inductively Coupled Mass Plasma Spectrometry (ICP-MS; PerkinElmer NexION 2000, University of Nottingham). We monitored 20 elements (Na, Mg, P, S, K, Ca, Ti, Cr, Mn, Fe, Co, Ni, Cu, Zn, As, Se, Rb, Sr, Mo, Cd and Pb) in the soil extracts samples. Individual soil samples of genotyped individuals (80 samples in total) were dried at 60°C. Soil samples were sieved afterwards. Samples were prepared according to the protocol (see attached Protocols). The original data are attached as Data S1a. We quantified the elemental soil and leaf composition of the elements in samples from reciprocal transplant experiment by ICP OES spectrometer INTEGRA 6000 (GBC, Dandenong Australia). We monitored three elements that were identified as key elements differentiating S and N soils of the natural populations (Ca, Mg, and Ni) and decomposed samples prior the analysis. For details see attached Protocols. The original data are attached as Data S1b.

### Screening natural variation in the *TPC1* locus

To screen a broader set of *TPC1* genotypes in the relevant serpentine populations S3-S5, we Sanger-sequenced additional individuals sampled at the original sites of the focal S3, S4 and S5 populations (11, 17, and 12 individuals, respectively) exhibiting non-synonymous variation in the *TPC1* locus. For the amplification of the exon around the candidate site we used specifically designed primers (amplifying genomic region scaffold_6:23,043,556– 23,043,854 bp in the reference; forward primer: CAAATTCAACAGAAGTAAAATAGTGATGACG; reverse primer: CTATCTATTGTTCAACTTCAATGACTACCCC). The mix for PCR contained 0.3 μl ofl of forward and reverse primer each, 14.2 μl ofl of ddH2O, 0.2 μl ofl of MyTag DNA polymerase, 4 μl ofl of reaction buffer MyTag, and we added 1 μl ofl (10 ng) of DNA. The PCR amplification was conducted in a thermocycler (Eppendorf Mastercycler Pro) under the following conditions: 1 min of denaturation at 95 °C, followed by 35 cycles: 20 s at 95 °C, 25 s at 60 °C, 45 s at 72 °C, and a final extension for 5 min at 72 °C. Amplification products of high purity were sequenced at 3130xl Genetic Analyser (DNA laboratory of Faculty of Science, Charles University, Prague).

Then, we checked whether the candidate alleles, inferred as serpentine specific in our dataset, are also absent in a published broad non-serpentine sampling among outcrossing *Arabidopsis* species^8,33,94–99^. We downloaded all the available short-read genomic sequences published with the referred studies, called variants using the same approach as described above and checked the genotypes at the candidate site (630 residue in *A. arenosa* and 633 in *A. lyrata* and *A. halleri*). In total, we screened 1,724 alleles of *A. arenosa*, 178 alleles of *A. halleri*, and 224 alleles of *A. lyrata*, summary of the investigated populations is attached in Dataset S11.

To visually compare variation at the entire *TPC1* locus, we generated consensus sequences for the group of all five non-serpentine *A. arenosa* populations and for each separate serpentine population using the vcf with all variants in the region of scaffold_6:23,042,733-23,048,601 using bcftools (Fig S14). Sites were included in the consensus sequence if they had AF > 50%. As the original VCF contained only biallelic sites, an additional multiallelic VCF was also created using GATK and any variants with AF > 50% were manually added to the biallelic consensus sequence. The *A. lyrata* and *A. thaliana* non-serpentine sequence were assumed to match the corresponding reference for each species.

Finally, we screened the variation in the *TPC1* locus at deep phylogenetic scales. We generated multiple sequence alignments using Clustal-Omega^100^ from the available mRNA sequences from GenBank (*Emiliania huxleyi, Hordeum vulgare, Arabidopsis thaliana, Triticum aestivum, Salmo salar, Strongylocentrotus purpuratus, Homo sapien, Rattus norvegicus and Mus musculus*) that were complemented by the consensus *Arabidopsis* sequences described above. Alignments were manually refined and visualised in JalView^101^.

### Structural Homology Models

Structural homology models of dimeric *TPC1* were generated with Modeller version 9.24^102^ using two *A. thaliana* crystal structures (5E1J and 5DQQ^46,48^) as templates. The final model was determined by its discrete optimized protein energy (DOPE) score.

## Acknowledgements

This work was supported by the Czech Science Foundation (project 20-22783S to FK), Charles University (project Primus/SCI/35 to FK and GAUK 410120 to VK), the European Research Council (ERC) under the European Union’s Horizon 2020 research and innovation programme [grant number ERC-StG 679056 HOTSPOT to LY]. Additional support was provided by the long-term research development project No. RVO 67985939 of the Czech Academy of Sciences. Computational resources were provided by the CESNET LM2015042 and the CERIT Scientific Cloud LM2015085. The authors thank Lenka Flašková, Gabriela Šrámková, Anna Krejčová, and Mellieha Allen for help with laboratory work and results interpretation.

