## Supplementary information for "Parallel adaptation in autopolyploid *Arabidopsis arenosa* is dominated by repeated recruitment of shared alleles"

##### Supplementary Datasets

**Supplementary Dataset 1**: Sequence processing and quality assessment of each re-sequenced individual.

**Supplementary Dataset 2A-E**: Description of differentiation candidates (1% of empirical Fst outliers) for five pairs.

**Supplementary Dataset 3A-C**: 3A) Results from GO enrichment analysis of the union of all 2245 differentiation candidates. 3B) Fisher exact test on overlaps among differentiation candidates from five pairs. 3C) Results from GO enrichment analysis of 207 parallel differentiation candidates.

**Supplementary Dataset 4A-E**: Description of LFMM candidates (genes with at least one SNP associated with soil content of at least one element - Ca/Mg, Co, Mg and Ni) across all samples.

**Supplementary Dataset 5**: Fisher exact test on overlap between parallel differentiation candidates and LFMM candidates.

**Supplementary Dataset 6A-B**: 6A) Summary of 61 serpentine adaptation candidates with identification of the variation source in DMC. 6B) Results from GO enrichment analysis of 61 serpentine adaptation candidates.

**Supplementary Dataset 7**: Summary of TE variants.

**Supplementary Dataset 8A-F**: A-D) Description of differentiation TE candidates (1% empirical $F_{ST}$ outliers) for four pairs. E) Fisher exact test on overlaps among differentiation TE candidates from four pairs. F) Description of parallel differentiation TE candidates.

**Supplementary Dataset 9**: Results from GO enrichment analysis of differentiation TE candidate genes.

**Supplementary Dataset 10**: Summary of 246 serpentine adaptation candidates with identification of the variation source in DMC**.**

**Supplementary Dataset 11**: Summary of the variation in allele frequencies of amino acid substitution in three wild *Arabidopsis (A. arenosa, A. halleri, A. lyrata)*.

**Supplementary Dataset 12**: Examples of parameter files used in fastsimcoal2 simulations to test for parallel origin of serpentine populations.

##### Supplementary Data

**Supplementary Data 1**: 1a) Individual soil concentrations from resequenced individuals. 1b) Individual leaf and soil concentrations from individuals cultivated in reciprocal transplant experiment.

##### Protocols

Protocols: DNA extraction and assesment of leaf and soil elements concentrations.

### Supplementary Tables

**Table S1** Details on sampled populations.

| pop code | pop | ploidy | pop name | n ind ionomics /genome | bedrock | altitude | lat | lon | country |
|---|---|---|---|---|---|---|---|---|---|
| BOR | S1 | 4x | Borovsko | 8/8 | serpentine | 416 | 49.68381 | 15.133255 | CZ |
| STG | S2 | 4x | Steinegg | 8/7 | serpentine | 414 | 48.62993 | 15.54256 | AT |
| GUL | S3 | 4x | Gulsen | 8/8 | serpentine | 628 | 47.28167 | 14.92764 | AT |
| OPP | S4 | 4x | Oppenberg | 8/8 | serpentine | 1750 | 47.46403 | 14.23989 | AT |
| PER | S5 | 4x | Pernegg | 8/8 | serpentine | 540 | 47.35512 | 15.33697 | AT |
| VLA | N1 | 4x | Vlastejovice | 8/8 | siliceous | 345 | 49.73496 | 15.17484 | CZ |
| FUG | N2 | 4x | Fuglau | 8/8 | siliceous | 436 | 48.63149 | 15.55723 | AT |
| ING | N3 | 4x | Ingeringgraben | 8/8 | siliceous | 950 | 47.28405 | 14.68154 | AT |
| VOR | N4 | 4x | Vorberg | 8/8 | siliceous | 1010 | 47.49876 | 14.16964 | AT |
| HOC | N5 | 4x | Hochlantsch | 8/7 | siliceous | 545 | 47.37 | 15.38667 | AT |

**Table S2** Summary of datasets and filtrations for genomic analyses.

| Analysis | Dataset | Input filtrations |
|---|---|---|
| **Faststructure** | 4dg SNPs | MFFG 0.2, DP < 8, pruning over 1kb windows and 10kb distance between the windows, MAF < 0.05 |
| **PCA** | 4dg SNPs | MFFG 0.2 and DP < 8 |
| **Treemix** | 4dg SNPs | MFFG 0.2 and DP < 8 |
| **Pairwise Fst, nucleotide diversity, Tajima's D, Fastsimcoal** | 4dg, SNPs + invariants | MFFG 0.2 and DP < 8 at the population level |
| **Fst scans** | all SNPs | MFFG 0.2 and DP < 8 at the population level |
| **LFMM** | all SNPs | missing data filtered out, MAF < 0.05 |
| **Differentiation of regions containing TE variants** | all SNPs<br>and TE variants | MAF < 0.05, MFFG 0.2 and DP < 8 |
| **DMC** | all SNPs annotated | MFFG 0.2 and DP < 8 |
| **DMC (neutral data)** | 4dg SNPs | MFFG 0.2 and DP < 8 |

Number of sites before filtrations in brackets:
All SNPs (11,744,200): biallelic SNPs pre-filtered following GATK best parctices
4dg sites (4,720,044): fourfold-degenerate sites
All SNPs annotated (8,808,984): same as all sites with available annotation for SNPs retained from SnpEff.
MAF: minor allele frequency
MFFG: maximum fraction of filtered genotypes (reflecting missingness)
DP: read depth

**Table S3** Summary of SNP and TE variation for each population pair.

| pop | n SNPs[1] | pop. pair | n SNPs[1] | n SNPs[2] | n TEs[3] | n of 1kb outlier windows (SNPs) | n of 1kb outlier windows (TEs) |
|---|---|---|---|---|---|---|---|
| S1 | 395,760 | S1-N1 | 547,071 | 4,320,217 | 9,798 | 953 | 91 |
| N1 | 438,869 | | | | | | |
| S2 | 449,103 | S2-N2 | 561,047 | 4,458,385 | 10,197 | 954 | 94 |
| N2 | 432,398 | | | | | | |
| S3 | 406,035 | S3-N3 | 505,296 | 3,410,315 | 10,183 | 957 | 91 |
| N3 | 365,023 | | | | | | |
| S4 | 384,194 | S4-N4 | 487,623 | 3,842,682 | 10,112 | 935 | 93 |
| N4 | 389,774 | | | | | | |
| S5 | 418,341 | S5-N5 | 452,303 | 3,028,157 | - | 726 | - |
| N5 | 354,981 | | | | | | |

[1] 4dg SNPs MFFG 0.2 and DP < 8
[2] all SNPs MFFG 0.2 and DP < 8
[3] filtered TE variants

**Table S4** Summary of population means of selected environmental variables (Temp = annual mean temperature, Prec = annual precipitation (both from average for 1970-2000 and spatial resolution ~1km$^2$), Elev = elevation. For soil variables, min, **mean**, and max ppm concentrations are presented in separate lines for each population).

| pop | Temp [°C] | Prec [mm] | Elev [m] | Mg [ppm] | Ni [ppm] | Co [ppm] | Ca/Mg | Ca [ppm] | P [ppm] | S [ppm] | K [ppm] |
|---|---|---|---|---|---|---|---|---|---|---|---|
| S1 | 7.8 | 605.0 | 416.0 | 1938.2 | 62.4 | 2.9 | 0.3 | 825.4 | 30.3 | 34.6 | 97.6 |
| | | | | **2324.5** | **136.6** | **5.7** | **0.6** | **1447.5** | **81.7** | **82.8** | **213.0** |
| | | | | 3338.2 | 248.8 | 9.8 | 1.2 | 2473.4 | 168.8 | 149.2 | 354.1 |
| S2 | 8.2 | 694.0 | 414.0 | 2523.6 | 101.4 | 6.1 | 0.3 | 1036.5 | 39.2 | 48.1 | 207.6 |
| | | | | **3877.7** | **156.3** | **9.3** | **0.5** | **1730.9** | **67.2** | **116.9** | **294.6** |
| | | | | 4983.2 | 259.9 | 17.5 | 0.6 | 2208.3 | 128.8 | 251.9 | 398.9 |
| S3 | 6.8 | 983.0 | 628.0 | 662.9 | 18.9 | 2.3 | 0.5 | 332.1 | 73.7 | 24.0 | 52.8 |
| | | | | **2247.2** | **50.3** | **6.8** | **1.1** | **2437.9** | **213.4** | **88.7** | **614.3** |
| | | | | 4836.1 | 117.1 | 15.2 | 2.5 | 4801.6 | 358.0 | 226.1 | 1103.1 |
| S4 | 1.4 | 1503.0 | 1750.0 | 971.1 | 37.1 | 2.7 | 0.3 | 956.7 | 22.7 | 49.0 | 67.2 |
| | | | | **3289.5** | **99.3** | **7.5** | **0.7** | **1890.1** | **87.5** | **145.6** | **209.5** |
| | | | | 4989.1 | 141.2 | 13.6 | 1.3 | 4035.5 | 150.9 | 427.5 | 357.2 |
| S5 | 7.2 | 896.0 | 540.0 | 1115.4 | 19.3 | 1.9 | 0.5 | 920.3 | 19.4 | 42.1 | 118.5 |
| | | | | **2013.7** | **70.0** | **4.0** | **1.3** | **2498.5** | **62.7** | **99.3** | **248.9** |
| | | | | 3326.5 | 132.5 | 6.9 | 2.5 | 5492.2 | 138.2 | 389.6 | 452.7 |
| N1 | 8.2 | 581.0 | 345.0 | 143.6 | 4.7 | 0.2 | 2.9 | 601.4 | 22.5 | 44.7 | 62.7 |
| | | | | **246.1** | **7.1** | **0.4** | **3.5** | **846.2** | **48.2** | **64.4** | **165.9** |
| | | | | 347.7 | 12.2 | 0.7 | 4.2 | 1154.5 | 70.5 | 94.5 | 353.1 |
| N2 | 8.3 | 684.0 | 436.0 | 167.5 | 0.7 | 1.0 | 2.7 | 665.4 | 25.6 | 11.3 | 100.6 |
| | | | | **223.7** | **1.1** | **1.9** | **4.4** | **923.0** | **54.0** | **40.3** | **176.2** |
| | | | | 289.8 | 1.8 | 3.5 | 8.5 | 1425.3 | 71.8 | 78.6 | 291.1 |
| N3 | 4.3 | 1200.0 | 950.0 | 176.8 | 4.2 | 1.3 | 1.2 | 1043.1 | 68.0 | 67.2 | 145.2 |
| | | | | **771.8** | **21.1** | **3.6** | **3.3** | **2039.5** | **137.8** | **100.7** | **574.3** |
| | | | | 1443.2 | 40.4 | 7.5 | 6.1 | 5137.7 | 230.8 | 160.2 | 1237.6 |
| N4 | 5.3 | 1299.0 | 1010.0 | 29.3 | 0.2 | 0.2 | 12.7 | 704.6 | 8.7 | 8.4 | 25.8 |
| | | | | **41.8** | **0.5** | **0.5** | **21.8** | **835.4** | **22.4** | **28.7** | **94.8** |
| | | | | 86.5 | 0.8 | 0.7 | 26.4 | 1102.9 | 55.5 | 53.8 | 221.2 |
| N5 | 6.8 | 914.0 | 545.0 | 773.4 | 1.2 | 0.9 | 1.7 | 2889.9 | 37.8 | 70.8 | 310.7 |
| | | | | **2311.9** | **1.8** | **2.1** | **2.6** | **5335.2** | **175.0** | **156.4** | **792.9** |
| | | | | 3662.3 | 2.9 | 4.1 | 4.2 | 7508.3 | 355.2 | 428.7 | 1511.5 |

**Table S5** Parameters used in parallel selection modelling in DMC.

| parameter | parallel selection scenario | values used |
|---|---|---|
| selection coefficient | all scenarios | 0.0001, 0.001, 0.01, 0.05, 0.1, 0.2, 0.5 |
| migration rates | migration | 1e-07, 1e-06, 0.00001, 0.0001, 0.001, 0.01 |
| initial allele frequencies prior to selection | standing variation | 2.5e-06, 0.00001, 0.0001, 0.001, 0.01 |
| times for which allele was standing prior to onset of selection[1] | standing variation | 100, 1000, 5000, 10000, 30000, 50000, 100000 |

[1] This time range spans from the low CI of the youngest split between S and N populations inferred by our coalescent simulations to approximately three times of the origin of the autotetraploid cytotype as inferred for *A. arenosa* by previous range-wide studies[1,2].

We used effective population size $N_e = 100{,}000$ calculated from the mean nucleotide diversity of our populations 0.029 following the equation $N_e = \pi/8\mu$ ($\mu = 4.3e^{-8}$ from ref. [3]).

We used recombination rate of $3.7e^{-8}$ estimated for the reference species *A. lyrata* previously[4].

**Table S6** Summary of allele frequencies of non-synonymous substitutions in the *TPC1* locus in all *A. arenosa* populations resequenced in this study (separately for each S pop., allele number (AN) = 28-32, and summed across all N pops, AN = 156). The high-frequency serpentine-specific alleles are highlighted in bold.

| Position | Substitution | S1 | S2 | S3 | S4 | S5 | N |
|---|---|---|---|---|---|---|---|
| scaffold_6:23047864 | Phe79Leu | 0.000 | 0.594 | 0.781 | 0.719 | 0.844 | 0.526 |
| scaffold_6:23047750 | Gln85Lys | 0.000 | 0.000 | 0.000 | 0.000 | 0.781 | 0.295 |
| scaffold_6:23047666 | Asn113Asp | 0.000 | 0.000 | 0.000 | 0.000 | 0.594 | 0.205 |
| scaffold_6:23047662 | Val114Ala | 0.000 | 0.000 | 0.000 | 0.000 | 0.656 | 0.314 |
| scaffold_6:23047015 | Leu190Ile | 0.906 | 0.875 | 1.000 | 0.938 | 1.000 | 0.929 |
| scaffold_6:23046782 | His200Asp | 0.563 | 0.688 | 0.969 | 0.688 | 1.000 | 0.750 |
| scaffold_6:23046751 | Gly210Val | 0.563 | 0.688 | 0.969 | 0.719 | 1.000 | 0.737 |
| scaffold_6:23046295 | Ile300Val | 0.656 | 0.813 | 1.000 | 1.000 | 1.000 | 0.853 |
| scaffold_6:23046128 | Arg320Lys | 0.594 | 0.750 | 0.969 | 0.781 | 1.000 | 0.769 |
| scaffold_6:23045888 | Glu349Gln | 0.594 | 0.688 | 0.969 | 0.781 | 1.000 | 0.744 |
| scaffold_6:23045881 | Asn352Thr | 0.688 | 0.781 | 1.000 | 1.000 | 1.000 | 0.840 |
| scaffold_6:23045354 | Lys409Gln | 0.594 | 0.688 | 0.969 | 0.688 | 1.000 | 0.756 |
| scaffold_6:23045270 | Val437Ile | 0.656 | 0.844 | 1.000 | 1.000 | 1.000 | 0.846 |
| scaffold_6:23044774 | Ala492Ser | 0.594 | 0.000 | 0.906 | 0.000 | 0.000 | 0.622 |
| scaffold_6:23044106 | Ile582Val | 0.719 | 0.813 | 1.000 | 1.000 | 0.906 | 0.853 |
| scaffold_6:23044055 | Leu599Met | 0.594 | 0.563 | 0.969 | 0.750 | 0.781 | 0.692 |
| scaffold_6:23043812 | Val630Ile | 0.000 | 0.000 | 0.031 | 0.000 | 0.031 | 0.090 |
| **scaffold_6:23043812** | **Val630Leu** | **0.000** | **0.000** | **0.938** | **0.000** | **0.469** | **0.000** |
| **scaffold_6:23043812** | **Val630Tyr** | **0.000** | **0.000** | **0.000** | **0.659** | **0.000** | **0.000** |
| scaffold_6:23043385 | Asn682Lys | 0.000 | 0.719 | 0.969 | 0.750 | 0.938 | 0.756 |
| scaffold_6:23043108 | Thr700Ser | 0.000 | 0.000 | 0.719 | 0.000 | 0.656 | 0.487 |

**Table S7** Chi-squared test for rotamer positioning of 630Tyr TPC1 alllele in a homodimer or heterodimer. Position is referred to as either ‘in’ when the residue points towards the channel or ‘out’ when the residue points away from the channel.

| **Homodimer** | **Expected** | **Actual** |
|---|---|---|
| 1 in, 1 out | 33.3 | 35 |
| 2 in | 33.3 | 20 |
| 2 out | 33.3 | 55 |
| P-value | | 0.000058 |
| **Heterodimer** | | |
| In | 50 | 41 |
| Out | 50 | 59 |
| P-value | | 0.0718606 |
| **Homo** | | |
| 1 in, 1 out | 45 | 35 |
| 2 out | 45 | 55 |
| P-value | | 0.035015 |

### Supplementary Figures

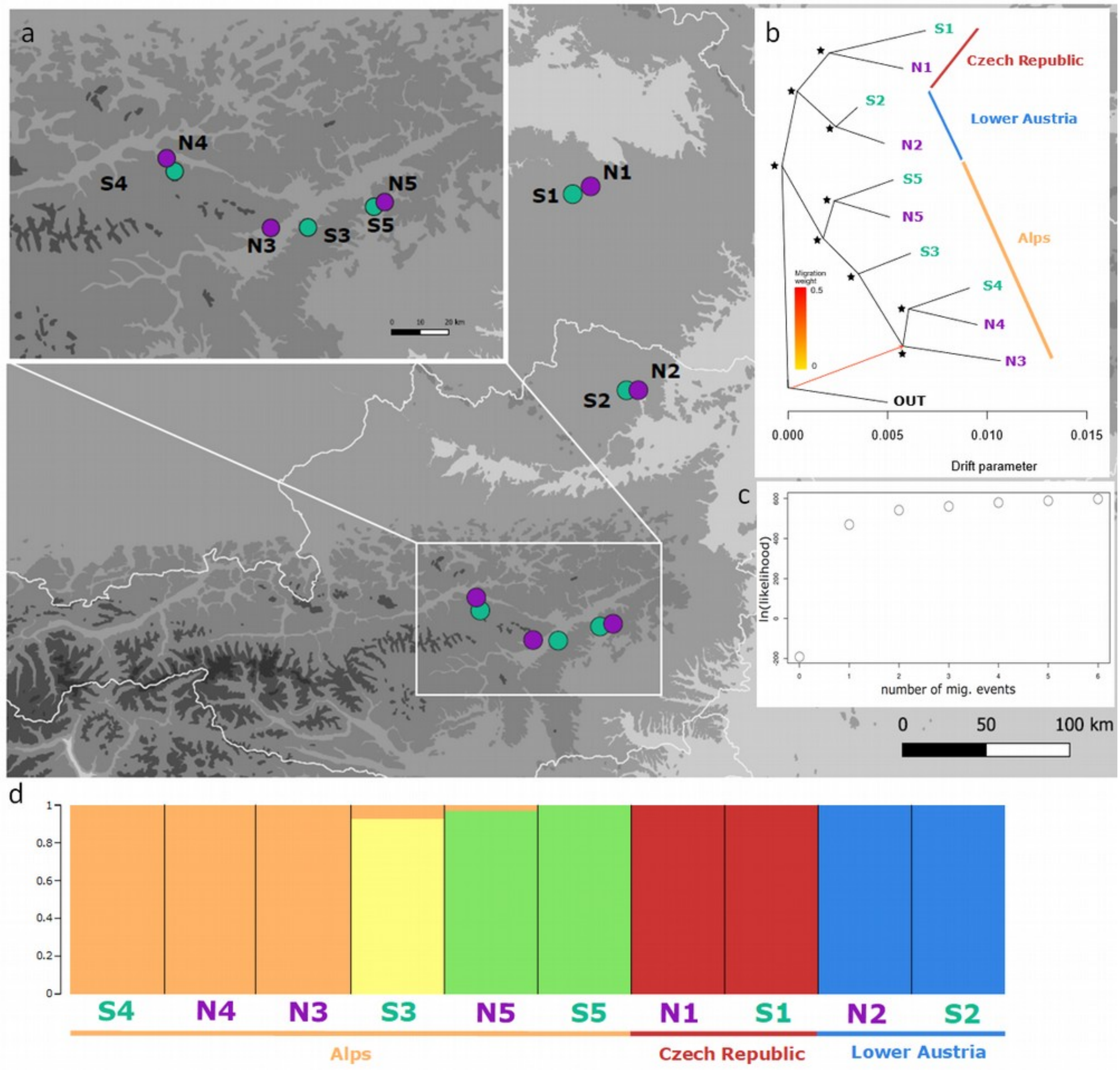


**Fig. S1** Geographic distribution and genetic structure of sampled autotetraploid *A. arenosa* populations in Central Europe inferred by analysis of 4dg SNPs. a) Locations of the populations. b) Allele frequency covariance graph of populations; asterisks show the 100 bootstrap branch support, red arrow shows one migration event. The outgroup (OUT) is represented by a tetraploid population from Western Carpathians, the ancestral area of tetraploid *A. arenosa* inferred previously[3]. c) Pattern of increasing likelihood with rising number of migration events from one to six in Treemix analyses, showing saturation after adding one migration event. d) Proportional assignment of individuals to five clusters (corresponding to the number of population pairs) inferred by fastStructure (9,923 LD-thinned 4dg SNPs). S – serpentine, N – non-serpentine.

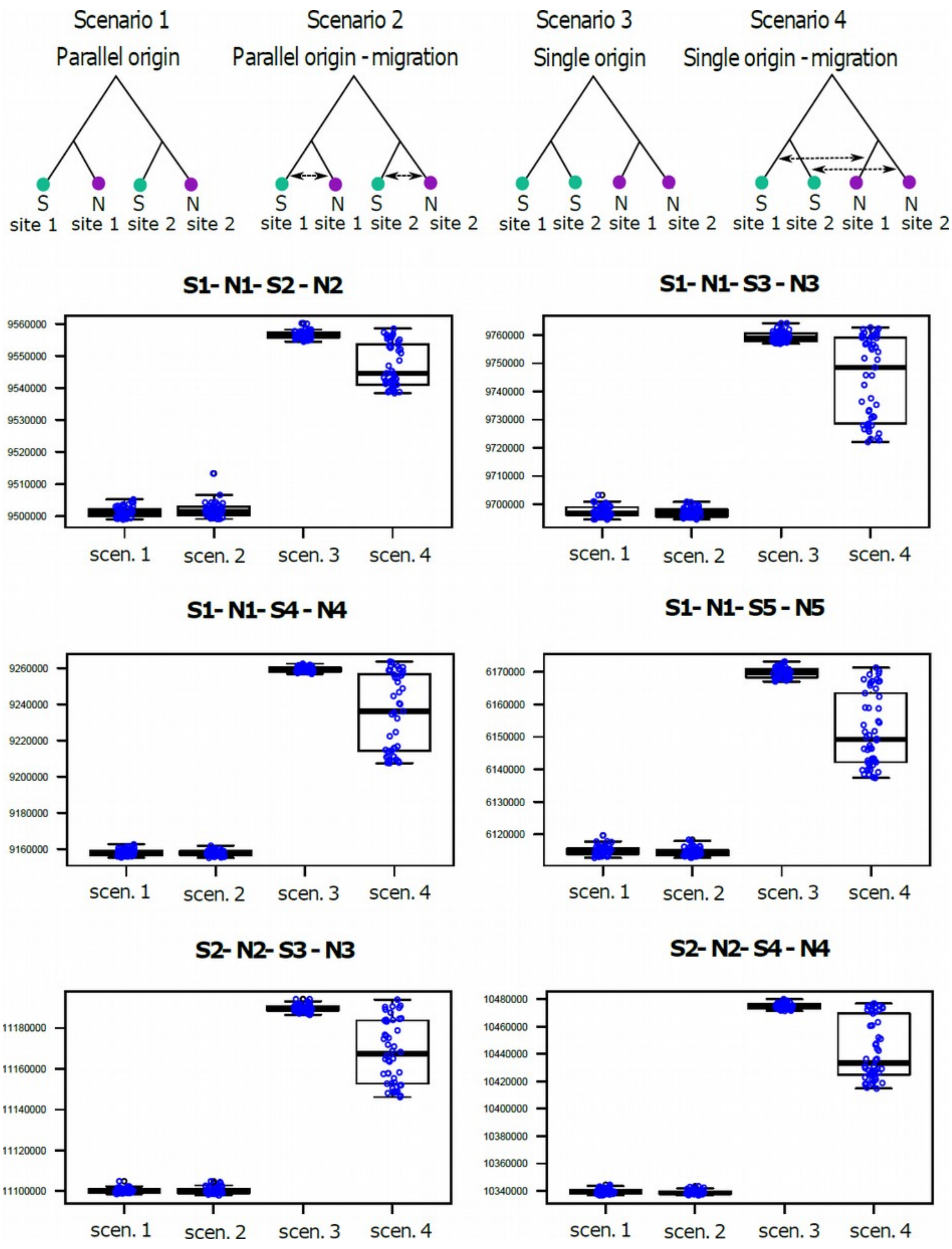
Scenario 1
Parallel origin
Scenario 2
Parallel origin - migration
Scenario 3
Single origin
Scenario 4
Single origin - migration
S site 1
N site 1
S site 2
N site 2
S1- N1- S2 - N2
S1- N1- S3 - N3
S1- N1- S4 - N4
S1- N1- S5 - N5
S2- N2- S3 - N3
S2- N2- S4 - N4
scen. 1
scen. 2
scen. 3
scen. 4

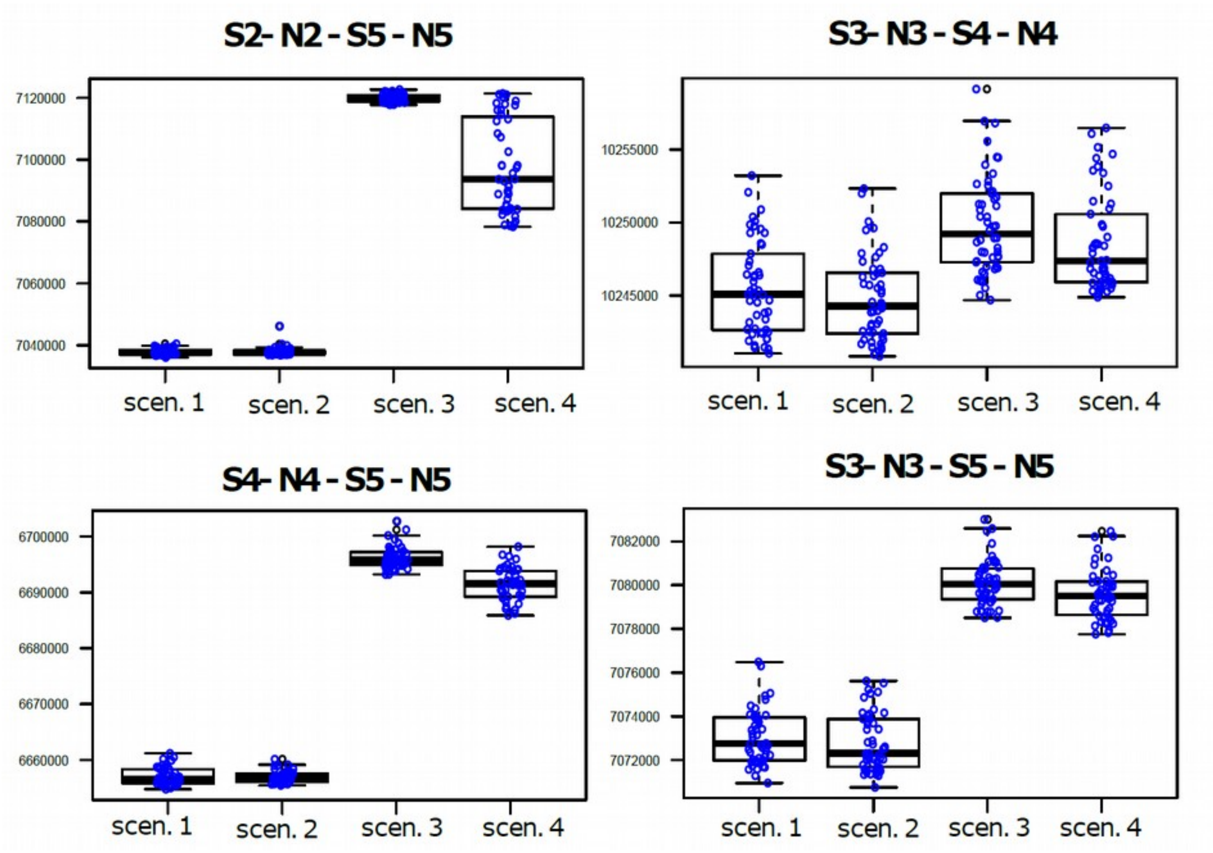


**Fig. S2** Comparison of Akaike information criteria (AIC) across four scenarios approximating the origin of serpentine populations in pairs of geographically proximal serpentine and non-serpentine populations. We iterated all pairs of serpentine (S) and non-serpentine (N) populations in a pairwise manner (i.e. 10 combinations). Each scenario was simulated by independent 50 fastsimcoal runs, the corresponding distribution of the AIC values over these 50 runs (blue dots) is summarized by the boxplots. The topologies of the evaluated scenarios are depicted on top. Consistently over all possible pairwise iterations the scenario of parallel colonisation of serpentine at each site was more likely than single origin of each edaphic type. Note that subsequent gene flow between substrate types within each S-N population pair was unlikely as the assumption of migration within each population pair had not significantly improved the model fit (scenario 1 vs. scenario 2 comparison).

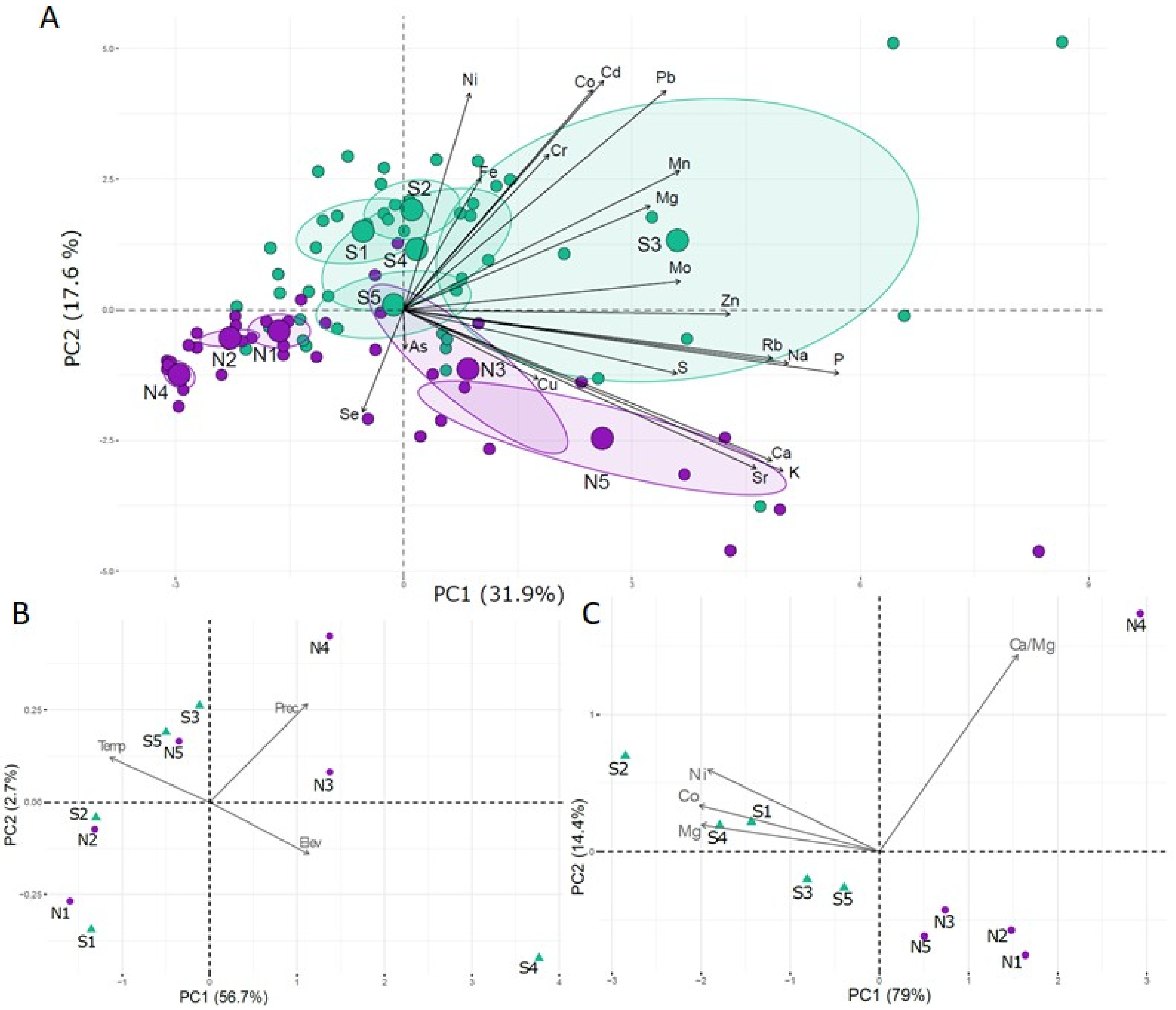


**Fig. S3** Environmental differentiation of the sampled serpentine (S, green dots) and non-serpentine (N, violet dots) sites displayed by unconstrained ordination (principal component analysis, PCA). a) Overall soil differentiation shown by PCA based on standardized concentrations of 20 elements recorded in rhizosphere soil of each sampled individual (95% confidence ellipses around population means) b) Lack of climatic differentiation between sister S and N populations shown by PCA based on three environmental variables (Temp = annual mean temperature, Prec = annual precipitation (both from average for 1970-2000 and spatial resolution ~1km$^2$), Elev = elevation. c) PCA based on the four soil variables significantly differentiating S and N soils (Ca/Mg, Co, Mg, and Ni) showing population means.

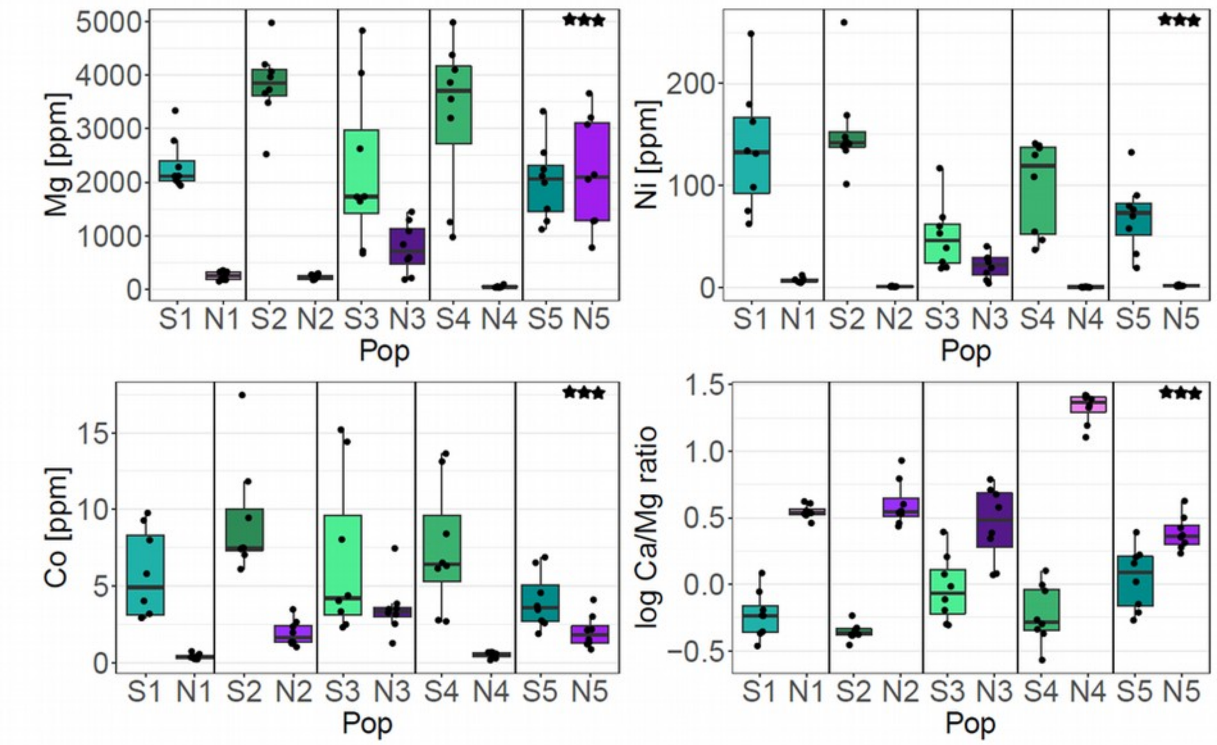


**Fig. S4** Variation in the concentrations of soil Mg, Ni, Co and Ca/Mg ratio (soil associated with each of the 80 sampled individuals). Note: Outlier S4_5 sample for Co concentration of 28.75 ppm was excluded to aid visibility. Differences between S and N soils in concentration of each element were tested by one-way ANOVA taking population pair as a random variable. Mg: $F_{1,77} = 71.7$, $p < 0.001$, Ni: $F_{1,77} = 117.4$, $p < 0.001$, Co: $F_{1,77} = 54.3$, $p < 0.001$, and Ca/Mg ratio: $F_{1,77} = 26.5$, $p < 0.001$.

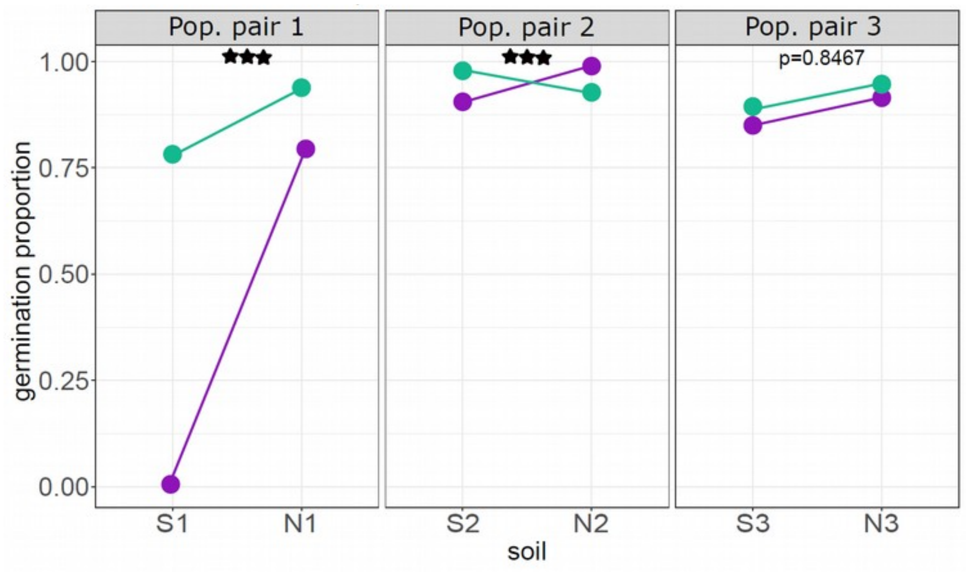


**Fig. S5** Differences in germination proportion in the originally serpentine (S, green) and non-serpentine (N, violet) populations transplanted to their native and alternative soil. Dots represent mean values across ~ 11 replicates (seed families) per treatment*population. Significances of the soil_treatment * soil_origin interaction terms were estimated by GLM with binomial errors, S1-N1 pair ($X^2 = 57.764$, $p < 0.001$), S2-N2 pair ($X^2 = 18.4083$, $p < 0.001$), and S3-N3 pair ($X^2 = 0.0374$, $p = 0.8467$).

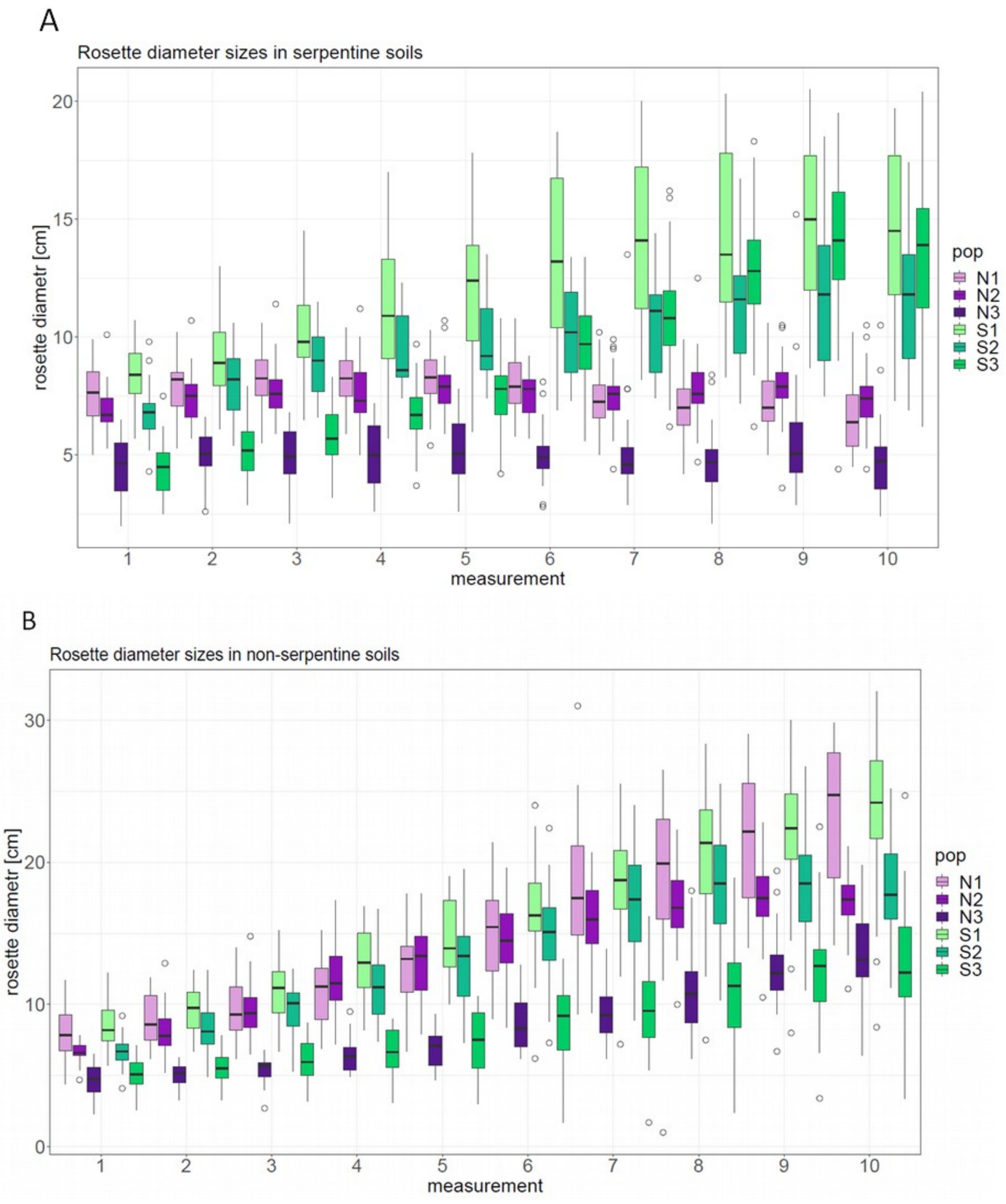


**Fig. S6** Differences in rosette diameter sizes of the individuals from the originally serpentine (S, green) and non-serpentine (N, violet) populations cultivated in serpentine (a) and non-serpentine soils (b). Data from three population pairs showing entire five-week period of cultivation. Note: rosettes were measured twice a week until all populations reached a plateau in the growth; ~ 25 replicates per population and treatment were cultivated. There was no mortality during the cultivation.

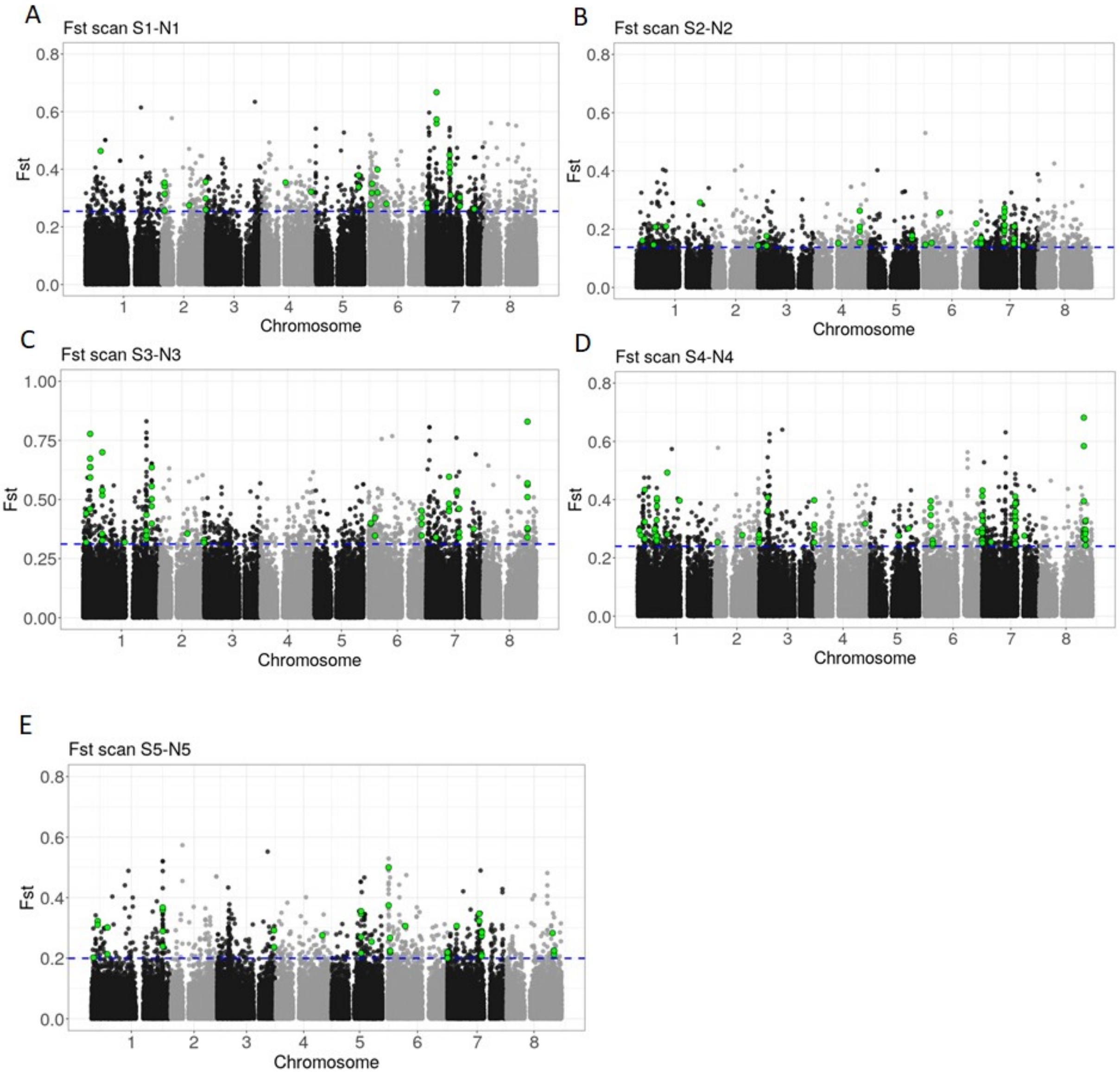


**Fig. S7** Window-based genomic differentiation between serpentine (S) and non-serpentine (N) population in each population pair (1–5, a–e, respectively) quantified by $F_{ST}$[5] calculated within 1 kb windows (dots) spanning the genome. Blue dashed line shows the upper 1% quantile, and green dots highlight position of the windows encompassing the 61 serpentine adaptation candidate genes (inferred downstream, see the main text).

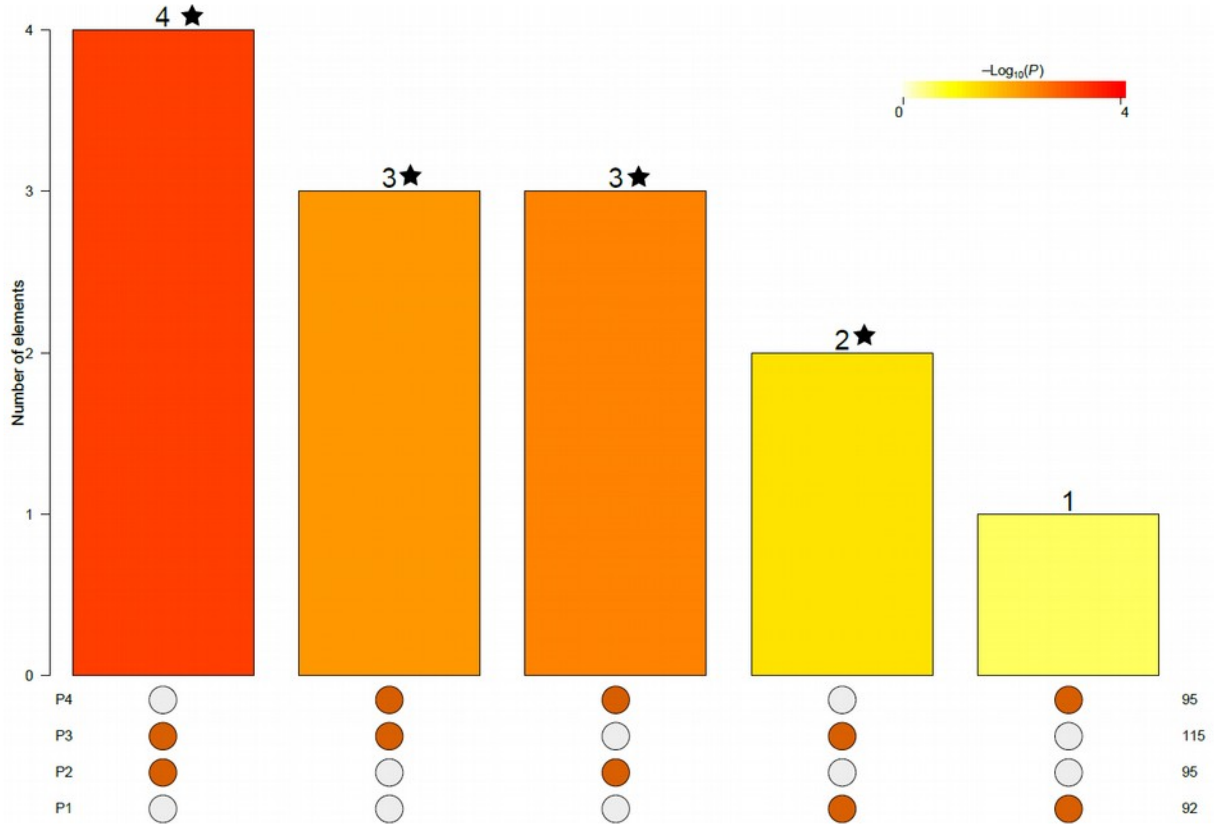


**Fig. S8** Intersection among lists of differentiation TE candidates (1% empirical $F_{ST}$ outliers) within each population pair (P1-P4); significant ($p < 0.05$) overlaps are marked with asterisks; the results of Fisher's exact test are available in Supplementary Dataset 8.

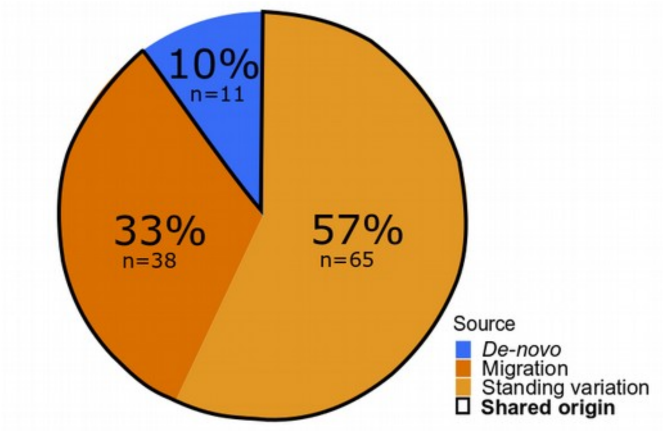


**Fig. S9** Sources of variation inferred by DMC analysis of the 246 serpentine adaptation candidates that were identified by applying less stringent differentiation threshold (upper 3 % differentiation outlier treshold). The candidates were further overlapped across population pairs leading to 1179 parallel differentiation candidates and further by 2809 LFMM candidates. Pie chart summarizes the proportions of cases of parallel adaptation (114 significantly non-neutral cases in total) reflecting likely origin from from *de-novo* mutations, shared from standing variation and by gene flow. The statistics for individual candidate loci are summarized in Supplementary dataset 10.

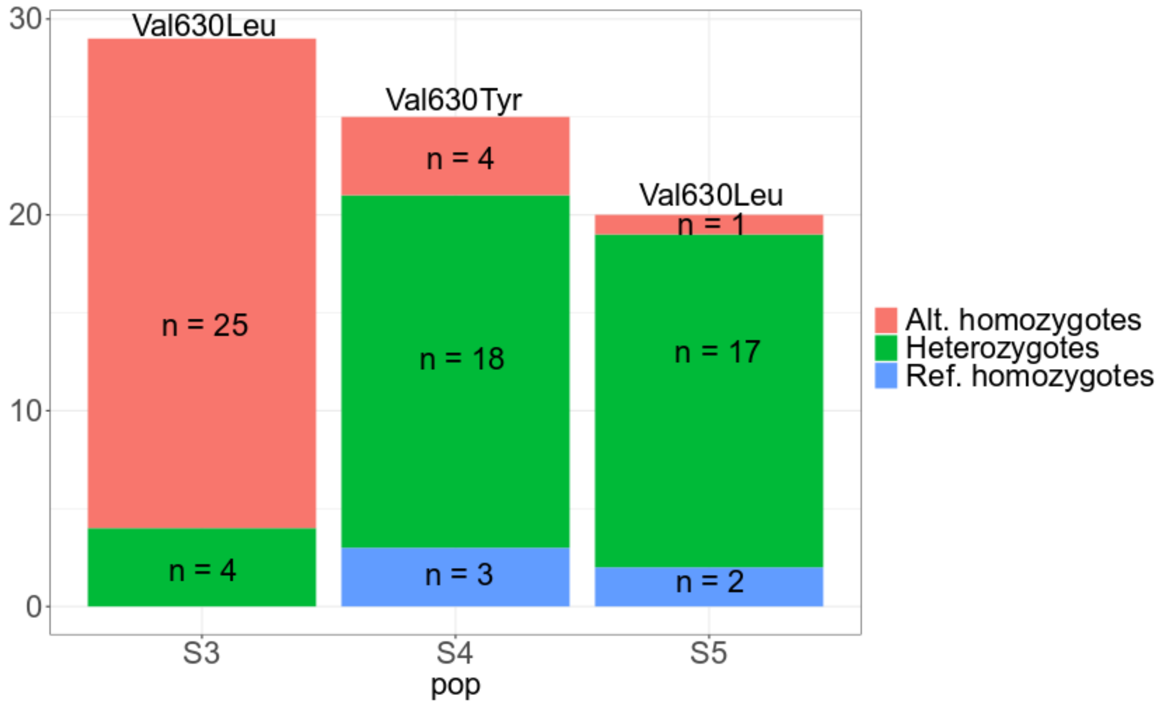


**Fig. S10** Genotype frequencies in residue 630 in the *TPC1* locus in the three serpentine *A. arenosa* populations encompassing serpentine-specific variants at this position. Alternative homozygous state (Alt. homozygotes) is represented by private allele C (mutation Val630Leu) in S3 and S5 populations and another private allele T in S4 population (mutation Val630Tyr). Heterozygous state is represented by alleles C/G in S3 population, A/C/G alleles in S5 population, and T/G in S4 population. Reference homozygous state (Ref. homozygotes) is represented by the widespread non-serpentine allele G (Val630Val).

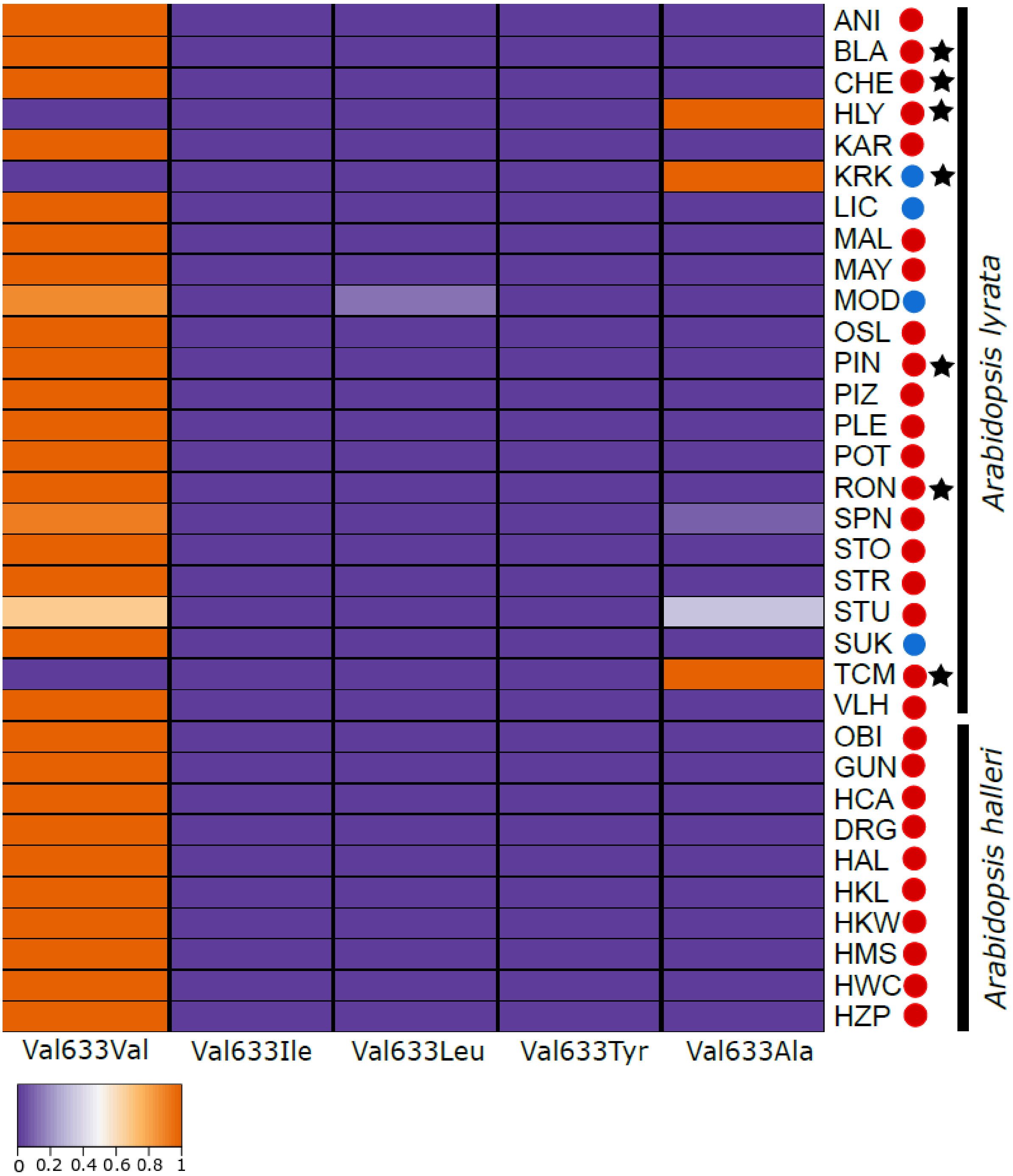


**Fig. S11** Allele frequencies of amino acid substitutions (colour key) in *A. lyrata* residue 633 (orthologous to 630 in *A. arenosa*) of the *TPC1* locus in *A. halleri* and *A. lyrata* populations inferred by a reanalysis of the available short read data. Red points: diploids, blue points: tetraploids. The substitution of Val633Leu found rarely in one *A. lyrata* population (MOD, limestone population) is encoded by a different codon (GTA -> TTA) than in *A. arenosa* serpentine populations (GTA -> CTA) and is thus non-homologous. “Population” samples represented by only one individual are marked by a star. On X-axis are the amino acid substitutions accompanied with the ancestral state Val633Val and on Y-axis are population codes for *A. halleri* and *A. lyrata* populations [6–12] (see Supplementary Dataset 11 for population details and original values).

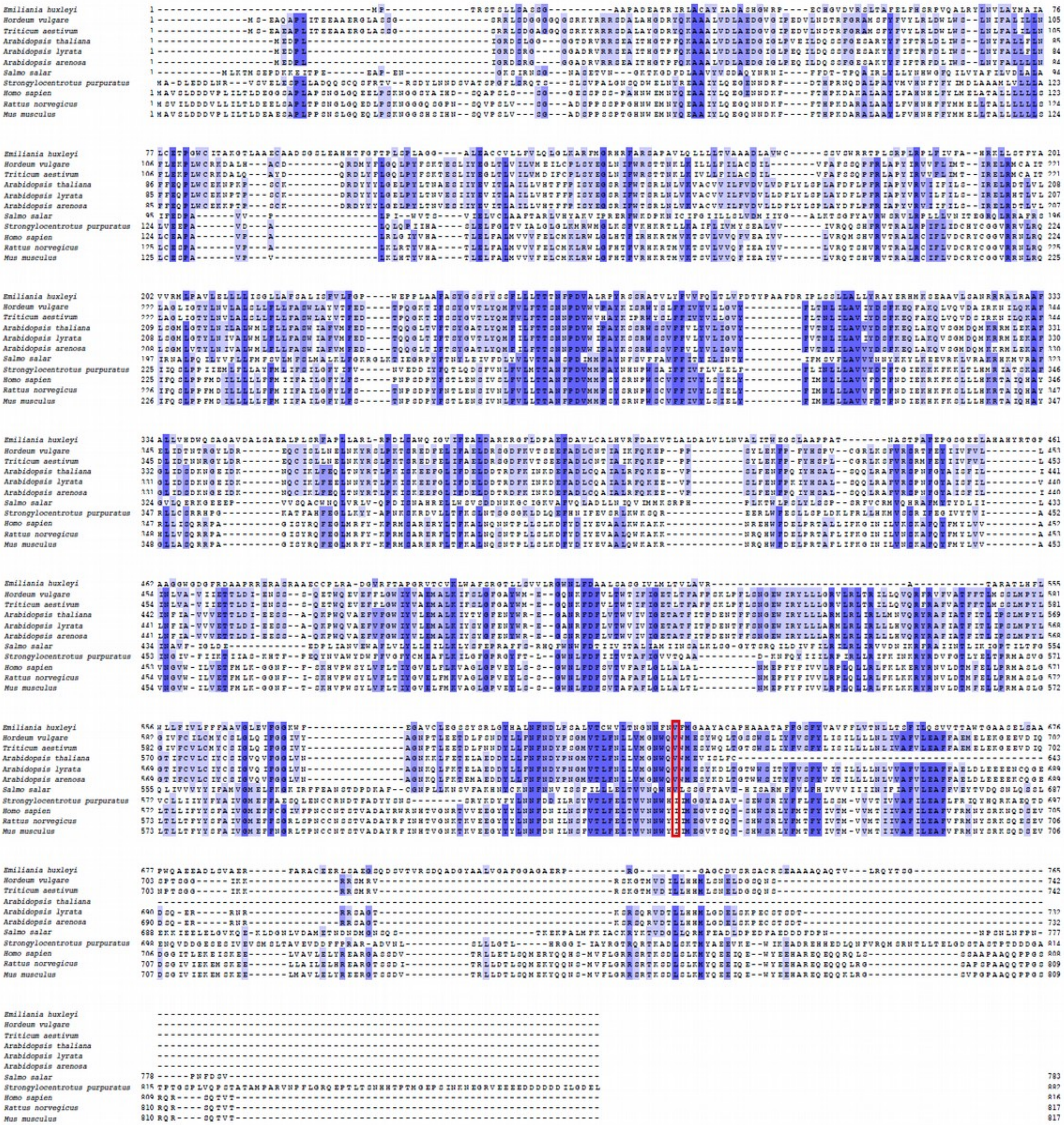


**Fig. S12** Cross-kingdom sequence conservation of the *TPC1* locus. Multiple sequence alignment of *TPC1* sequences (*Emiliania huxleyi, Hordeum vulgare, Arabidopsis thaliana, Triticum aestivum, Salmo salar, Strongylocentrotus purpuratus, Homo sapiens, Rattus norvegicus* and *Mus musculus*). Residues are coloured according to the percentage that match the consensus sequence from 100% (dark blue) to 0% (white). The position of the serpentine – specific high frequency non-synonymous SNP (Val634 in *A. thaliana*, Val630 in *A. arenosa* and Val633 in *A. lyrata*) is highlighted in red. Non-serpentine consensus sequences are shown for the three *Arabidopsis* species.

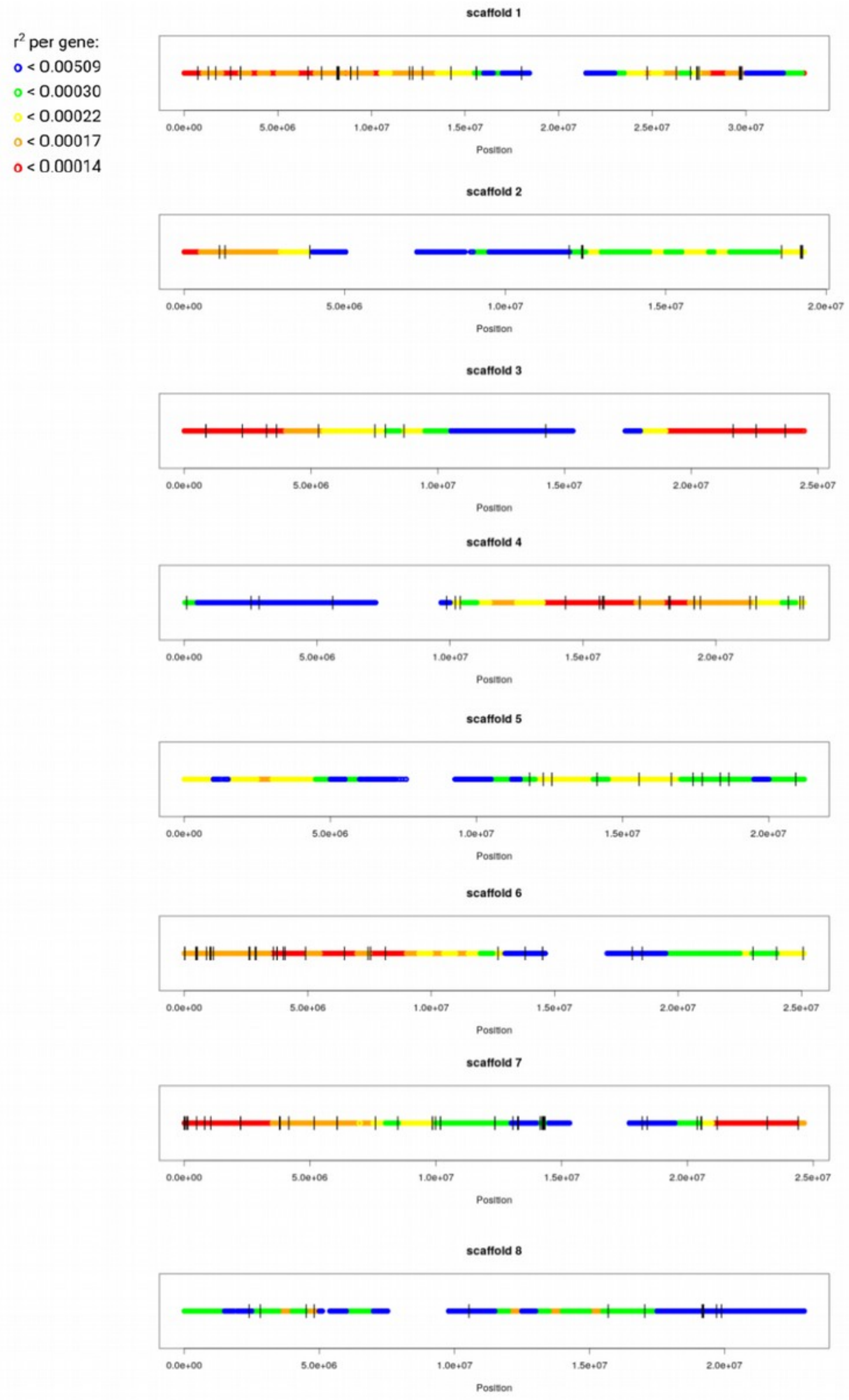


**Fig. S13** Location of all parallel Fst candidates (black vertical lines) on *A. lyrata* reference chromosomes colored by bins of distinct recombination rate per gene as estimated based on the available *A. lyrata* genetic map[4]. The figure illustrates that parallel Fst candidates are distributed throughout the genome, not being limited to regions with low recombination rate per gene (red/orange).

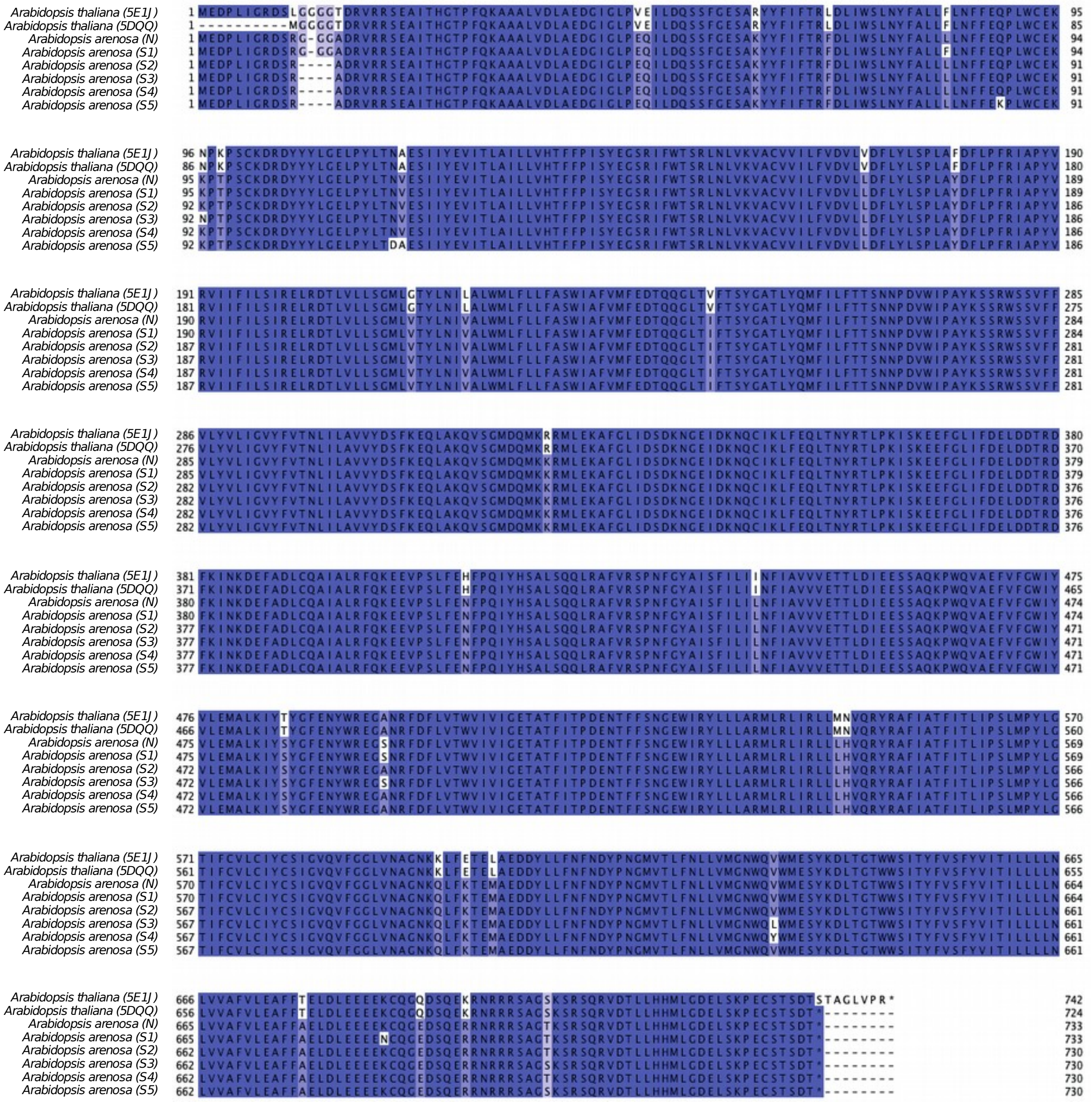

**Fig. S14** Multiple sequence alignment of *TPC1* consensus sequences used for structural homology modelling*.* Shown are consensus sequences from non-serpentine *A. arenosa* (N), five serpentine *A. arenosa,* and two non-serpentine *A. thaliana* crystal structures (5E1J and 5DQQ[13,14]). Residues are coloured according to percentage identity from 100% (dark blue) to 0% (white). Residue 630 in *A. arenosa* (634 in *A. thaliana* and 633 in *A. lyrata)* where the only serpentine-specific variation occurs is highlighted in red frame. The eight other nonsynonymous polymorphisms showing within *A. arenosa* variation are not serpentine specific (Table S6) and sit away from any known functional sites, indicating that they are likely non-functional (Fig 3d).
